# Characterization of Liquid-Liquid Phase Separation of Companion of Cellulose Synthases under Stress Mimicking Conditions

**DOI:** 10.64898/2026.09.07.749946

**Authors:** Viswanathan Gurumoorthy, Qiu Zhang, Wellington Leite, Alan C. Hicks, Jaydeep L. Kolape, Rajan Lamicchane, Hugh O’Neill

## Abstract

Companion of Cellulose Synthase (CC), has previously been shown to sustain cellulose synthesis during salt stress. However, the role of CCs in this process is not well-understood. In this study, we show that the intrinsically disordered N-terminal domain of Arabidopsis CC1 (CC1NTD) can undergo liquid-liquid phase separation (LLPS) under defined reducing conditions in the presence of trimethylamine N-oxide (TMAO), a naturally occurring plant stress osmolyte. The observed biomolecular condensates are enriched in CC1NTD and display liquid-like behavior, including spherical morphology, fusion, and dynamic exchange with the surrounding solution. We further show that TMAO compacts CC1NTD without inducing folding, that CC1NTD condensates recruit tubulin and accelerate microtubule polymerization while retaining dynamic properties. Additionally, we also studied the intrinsically disordered N-terminal domain of CC2, a closely related paralog to CC1. We found that CC2NTD remains disordered but does not phase separate independently under the same experimental conditions. However, it does partition into CC1NTD condensates. Together, these results define a condition-dependent condensate state for CC1NTD and link that state to tubulin-related function. Finally, we propose a novel mechanism for the formation and maintenance of stress-associated cellulose synthase compartments driven by LLPS of the disordered regions of CSC and accessory proteins.

## Introduction

Liquid-liquid phase separation (LLPS) is now recognized as a fundamental mechanism for the formation of membraneless organelles, also referred to as biomolecular condensates ^1^. Under defined physicochemical conditions, a homogeneous mixture of biomacromolecules such as proteins and nucleic acids can demix into two immiscible phases: a dense phase, often visible as liquid-like droplets, and a surrounding dilute phase. Biomolecular condensates are now known to participate in a wide range of cellular functions. They can sequester specific molecules, buffer the concentration of biomolecules, and locally enrich components required for biochemical reactions ^2^. Because phase separation is sensitive to physicochemical conditions, condensate formation can also serve as a means of responding to changes in the cellular environment. A growing number of biomolecular condensates have been described across the kingdoms of life, and plant condensates are increasingly recognized as important in processes including development, signaling, and stress responses ^3^. Some recent examples in plants include flowering time ^4,5^, immunity ^6,7^, hormone response ^8^, stress responses ^9,10^, embryo development ^11^, and seed germination ^12^.

Salt stress is a major agricultural challenge that disrupts plant growth and strongly affects cellulose synthesis and cortical microtubule organization. Adaption mechanisms to counter salt stress include plant hormone-mediated stress resistance signaling, reduced photosynthetic rate, modulation of reactive oxygen species metabolism and osmoregulation through small solutes to counteract increased osmotic pressure ^13–16^. Morphologically, cell walls have been shown to undergo remodeling and are believed to act as early sensors of stress and control plant cell division and growth under stress ^16,17^. Similarly, depolymerization and reorganization of plant cortical microtubule arrays have been found to be an important adaption mechanism for salt stress tolerance ^18–20^.

Arabidopsis Companion of Cellulose Synthase (CC) family proteins have been identified as important for sustaining cellulose synthesis during salt stress ^18^. Knockout mutants lacking CC1 and CC2 show reduced hypocotyl growth and impaired cellulose synthesis under salt stress, and CC1 has been linked to both cellulose synthase complexes and cortical microtubule organization ^18^. These observations suggest that CC proteins are part of the machinery that helps maintain cell wall synthesis and cytoskeletal organization under stress. CC1 protein contains three domains: an N-terminal intrinsically disordered cytosolic domain, a transmembrane helix, and a C-terminal β-sheet-rich domain that is predicted to reside in the apoplast ^18,21^. The N-terminal domain of CC1 (CC1NTD) is of particular interest because it is sufficient to rescue major stress-related defects in *cc1cc2* knockdown plants and to prevent disruption of cortical microtubule arrays ^18^. CC1NTD also interacts directly with microtubules, as shown by in vitro spin-down assays, in vivo colocalization, nuclear magnetic resonance (NMR), and cross-linked mass spectrometry ^22^. Previous work from our group showed that CC1NTD adopts an extended conformation during microtubule interaction, consistent with a role in bridging or bundling microtubules ^21^. These properties make CC1NTD a useful system for examining how a plant intrinsically disordered, microtubule-associated domain responds to stress-mimetic conditions.

Under salinity stress, plants accumulate osmolytes that help counter osmotic imbalance and influence protein stability and function ^23^. One such molecule is trimethylamine N-oxide (TMAO), which has been identified in plants under salt and drought stress and increases with prolonged stress exposure. An RNA-sequencing and mass spectrometry study identified accumulation of TMAO in plants during salt stress and drought stress ^24^. TMAO is also known as an osmolyte that can alter protein conformational ensembles and stability in vitro ^25–28^. Because CC1NTD is an intrinsically disordered cytosolic domain, TMAO provides a useful defined perturbation for testing how stress-associated osmolyte conditions affect its conformation and assembly behavior. In this study, TMAO is therefore used as a stress-mimetic condition to probe the biophysical response of CC1NTD.

During the course of our work on CC1NTD-microtubule interactions ^21^, we observed that concentrated CC1NTD solutions became cloudy and contained droplet-like assemblies, suggesting that CC1NTD can undergo LLPS under certain conditions. This observation motivated us to test its phase behavior under defined reducing, TMAO-containing conditions. Because a closely related paralog, CC2, also contains an intrinsically disordered N-terminal domain, comparison of CC1NTD and CC2NTD provides an opportunity to identify features that are specifically associated with condensate formation rather than with intrinsic disorder alone. Here, we examine whether TMAO-containing conditions promote condensate formation by CC1NTD, how these conditions alter the conformation of CC1NTD and whether CC1NTD condensates remain functionally active toward tubulin interactions and microtubule assembly. We further compare this behavior with that of CC2NTD. By focusing on these questions, this study defines the condensate behavior of CC1NTD under stress-mimetic conditions and tests how condensation is linked to conformational remodeling and microtubule-related activity.

## Materials and Methods

### Protein Sequence Analysis

Protein sequence alignment was performed using MUSCLE multiple sequence alignment v. 3.8.425 ^29^. IUPred was used to predict disordered regions ^30^. Prediction of liquid-liquid phase separation was performed using FuzDrop ^31,32^, catGRANULE ^33^, and PSPredictor ^34^.

### Protein Overexpression and Purification

The synthetic genes encoding CC1NTD (1-120 aa) and CC2NTD (1-98 aa) N-terminal domains were cloned into pET28a+ plasmid that has a hexahistidine tag at the C-terminal end. The purification procedure for protiated and deuterated CC1NTD was reported previously ^21^ and the CC2NTD purification procedure followed a similar approach. The details of the expression and purification of CC2NTD are described in Supplementary Information (Section 1). The method for fluorescent labeling of the amino terminal NH_2_ group of both CC1NTD and CC2NTD using Alexa Fluor 488 NHS succinimidyl ester (Invitrogen, USA) for fluorescent imaging and assays is described in the Supplementary Information (Section S2).

### Turbidity measurements

Protein turbidity (100µL) was measured using a SpectraMax (Molecular Devices, USA) spectrophotometer at 340nm ^35^ in a 96-well clear bottom plate (CellBIND™ Corning, USA) at 20s intervals for 30min at 20°C with orbital shaking for 3s between the reads. The protein concentration was varied between 0 – 60μM or 0 – 600μM for reducing and non-reducing conditions, respectively, in 50mM Tris-HCl, 150mM NaCl, pH 8.0 with or without 2mM (tris(2-carboxyethyl)phosphine) (TCEP) as a reducing agent (control buffer). The same buffer with varying amounts of TMAO (up to 3M) was termed the osmolyte buffer. The appropriate background and control measurements were also measured. Data were presented as mean ± standard error of the mean (sem).

Supernatant depletion experiments were performed to quantify how much protein partitioned to the dense and dilute phases. The turbid solutions were centrifuged at 14,000 x g for 5min and the protein concentration in the supernatant was determined using the Pierce™ 660nm protein assay. This was performed following the manufacturer’s protocol using lysozyme as a standard.

### Laser scanning confocal microscopy with differential interference contrast (DIC)

Condensates of CC1NTD and CC2NTD individually, CC1NTD-CC2NTD co-condensates, tubulin-CC1NTD co-condensates, and microtubule polymerization in the presence of CC1NTD condensates were imaged using confocal microscopy (Leica SP8 white light laser system, Leica Microsystems, Germany) equipped with a 63× oil objective lens. The details of sample preparation can be found in the Supplementary Information (Section S3). The apparent partition coefficient of proteins inside condensates was calculated as the ratio of fluorescence intensity inside condensates to outside (immediate background). Fluorescence intensity and size of droplets were quantified using ImageJ ^36^ from ≥10 droplets across three replicates. The data were presented as mean ± sem.

### Circular dichroism spectroscopy

Circular dichroism (CD) spectroscopy measurements for CC1NTD and CC2NTD were obtained using a Jasco J-810 CD spectrometer. Data were collected at room temperature in 0.1mm path length quartz cuvettes. Spectra were recorded every 0.1nm from 250 to 180nm. Due to strong absorption by the buffer below 195nm, the data below that value were excluded from analysis. Secondary structure content was estimated using BestSel online server (https://bestsel.elte.hu/) ^37^.

### Small-angle X-ray scattering

Small-angle X-ray scattering (SAXS) measurements were carried out using the Rigaku BioSAXS-2000 (Rigaku, Tokyo, Japan). using a configuration with a minimum Q of ∼0.008 Å^-1^. In total, 24 datasets were collected over 240min of exposure of each sample with an image refresh rate of 10min per image. The raw data were reduced to 1D data using the SAXSLab software package (Rigaku). Data analysis for model-independent approaches was done using BioXTAS RAW software package ^38^. Model-dependent analysis was done using SasView application package ^39^. Debye Gaussian coil model was fit using the following form factor (P(Q)) equation:

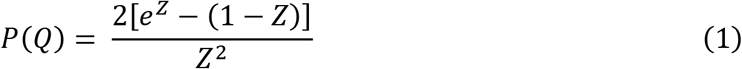

where 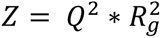 and the average persistence length (*l_p_*) can be calculated as follows ^40^:

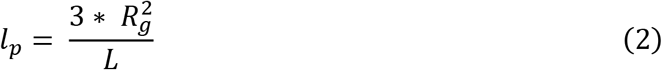

where *L*, the polypeptide chain length, is the number of amino acids multiplied by 3.63Å, the average amino acid length (*l*) ^41^. In the case of CC1NTD (135 amino acids including the C-terminal hexahistidine tag), this value is 490 Å. The average number of amino acids *n_a_* per persistence length is given by 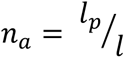^42^. GASBOR, part of the ATSAS suite, was used for ab initio reconstruction of CC1NTD models ^43^.

### Microtubule polymerization assay

Tubulin polymerization assay was performed according to the manufacturers protocol (Cytoskeleton Inc, BK006). Briefly, a 96-well plate was pre-warmed at 37°C for 30min. CC1NTD or CC2NTD (∼3mg/mL, ∼200µM) was diluted in either ice-cold general tubulin buffer or 1.5 M TMAO buffer, followed by addition of tubulin buffer (kept at 37°C) to each well. Buffer-only conditions (general tubulin buffer or TMAO buffer) were included as controls. Tubulin, maintained on ice, was rapidly added to each well to initiate polymerization, and measurements were started immediately. Microtubule polymerization was monitored by measuring absorbance at 340 nm using a SpectraMax plate reader (Molecular Devices, USA) at 37°C for 45min, with data acquisition every 30s. Absorbance values were corrected for background to yield ΔA at 340nm. The total polymer mass was calculated by averaging the last three timepoints of plateau phase as described previously ^18^. The data was presented as mean ± sem. The polymerization curve was fitted using the following Boltzmann equation:

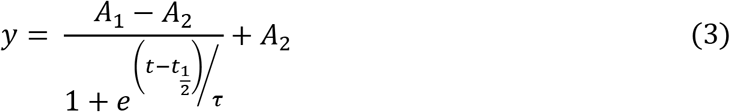

where *A*_1_ and *A*_2_ are lower and upper asymptotes; 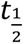 is half-time; and *τ* is the characteristic time scale that describes the sharpness of the polymerization transition, with smaller values corresponding to a steeper rise ^44–46^.

### Dynamic light scattering measurements

CC1NTD solutions and osmolyte buffer were first filtered using 0.5mL VWR® (0.2µm) centrifugal filters by centrifuging at 12,500 x g for 5min to remove any impurities. CC1NTD was diluted using either the control buffer or osmolyte buffer to achieve a 60µM CC1NTD in 1.5M TMAO solution. For tubulin experiments, 50µL of ice-cold tubulin (90µM) in polymerization buffer (80mM piperazine-N,N′-bis(2-ethanesulfonic acid), 2mM MgCl_2_, 0.5mM EGTA, 10% glycerol, 2mM GTP, pH 6.9) was quickly added to CC1NTD condensates in the osmolyte buffer and measured at 20 °C. The hydrodynamic radius was calculated using a Wyatt DynaPro Nanostar dynamic light scattering instrument (Wyatt Technology, USA). Details of this analysis are in the Supplementary Information (Section S4).

### Fluorescence recovery assay

For fluorescence recovery after photobleaching (FRAP) measurements, the CC1NTD condensates or tubulin–CC1NTD co-condensates were prepared in same way as described for microscopy measurements described above. The condensates were imaged before and directly after bleaching (circular regions of interest (ROIs); diameter ∼2–2.5µm) with a 488nm laser (90% intensity; 3 loops), and the fluorescence recovery of the bleached region was measured for 360s. For background, similar ROI was recorded outside the droplet region in parallel to measurements. For all measurements, the FRAP curves were background corrected and normalized. Experiments were performed using Leica confocal microscope (Leica SP8 white light laser system, Leica Microsystems, Germany) equipped with 63× oil objective. Experiments were repeated three times, and intensity values were averaged. Details of FRAP data fitting and analysis can be found in the Supplementary Information (Section S5).

### Small-angle neutron scattering (SANS)

SANS measurements were performed using the Bio-SANS instrument at the High Flux Isotope Reactor, Oak Ridge National Laboratory ^47^. Two configurations were used for the experiment. For the long configuration, the main detector was positioned at 15.5 from the sample, the wing detector was located at 1.13m and rotated by 5°, and the mid-range detector was located at 4m and rotated by −1°. Measurements used neutron wavelengths of 6 and 18Å, providing a combined *Q-*range of 0.001 – 0.8Å^-1^. Protiated CC1-NTD–tubulin and deuterated CC1-NTD– tubulin in 85% D₂O were also measured using a shorter sample-to-detector configuration with four neutron guides. For these measurements, the main detector was positioned at 7m, the wing detector at 1.13m and rotated 3.2°, and the mid-range detector at 4m and −2.7°. A neutron wavelength of 6Å provided a *Q*-range of 0.007 – 0.8Å^-1^. The momentum transfer was defined as *Q* = 4*πsin*(*θ*)/*λ*, where 2*θ* is the scattering angle and λ is the neutron wavelength. The instrument operated for both configurations with *Δλ*/*λ* = 13.2%. The 2D SANS data were corrected for instrument background, detector sensitivity, and instrument geometry, circularly averaged, and reduced to 1D scattering profiles using the facility data-reduction software, drt-SANS. Reference measurements, including the direct beam and corresponding buffer blanks, were collected for data reduction and intensity normalization. The buffer scattering was subtracted from the sample scattering to obtain the final background-corrected profiles.

Samples contained 60µM (0.89mg/ml) CC1NTD in 50mM Tris-HCl pH 8.0, 1.5M TMAO, 2mM TCEP, and 150mM NaCl. Either protiated or deuterated CC1NTD was used depending on the desired contrast condition. Protiated porcine tubulin (10mg/ml) (Cytoskeleton Inc.) was prepared in 80mM piperazine buffer, pH 6.9, containing 1mM GTP in D_2_O and was added to a final concentration of 2.5mg/ml (45µM) to the CC1 components directly before the SANS measurement. The final solvent contained either 42% or 85% D_2_O, depending on the contrast condition. Measurements were performed in 1 or 2mm path-length cylindrical quartz cuvettes (Hellma) at 20°C.

### Small-angle neutron scattering data analysis

The SANS profiles were analyzed using a combined Unified fit ^48,49^ and core–shell cylinder model. The Unified Fit contribution describes the characteristic size and scaling behavior of mesoscale organization within the condensates, whereas the core–shell cylinder term describes the cross-section of smaller elongated assemblies that are visible in solutions containing tubulin. The total scattering intensity was therefore expressed as the sum of the two contributions. The total scattering intensity was expressed as:

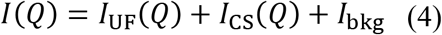

where *I*_UF_(*Q*) is the Unified Fit contribution, *I*_CS_(*Q*) is the core–shell cylinder contribution, and *I*_bkg_ is a *Q*-independent background. Additional details on the fitting approach and equations are provided in the Supplementary Information (Section S7).

## Results

### CC1NTD forms condensates in the presence of salt stress osmolyte TMAO

In previous work, we reported on the solution properties of recombinantly expressed CC1NTD using SAXS, Single-molecule Förster Resonance Energy Transfer, and computational modeling ^21^. CC1NTD was initially observed to form turbid solutions under non-reducing conditions at high protein concentration (∼400µM, 6mg/ml), and DIC imaging of these samples revealed spherical droplets that were approximately 1.5 ± 0.1 µm in diameter (Fig. S1a,b). Sequence-based predictors were consistent with an intrinsic propensity of CC1NTD to undergo phase separation. FuzDrop ^31^, PSPredictor ^34^, and catGRANULE ^33^ all showed positive scores for phase separation (Fig. S1d) and FuzDrop identified that CC1NTD has four aggregation-prone regions and a high droplet forming propensity (Fig. S1c,d). This observation motivated us to examine CC1NTD phase behavior under defined conditions including a reducing buffer environment (2mM TCEP) to mimic intracellular conditions and with trimethylamine N-oxide (TMAO), an osmolyte that is produced under salt stress conditions in plant cells.

Turbidity assays were performed to determine the conditions under which CC1NTD forms condensates. Protein concentration was varied from 20 to 60µM and TMAO concentration from 0 to 1.5M. CC1NTD remained non-turbid at lower TMAO concentrations, whereas turbidity became detectable at TMAO concentrations of approximately 1.0M and above. Increasing TMAO shifted the onset of turbidity to lower CC1NTD concentrations, indicating that condensate formation depends on both protein and osmolyte concentration (Fig. 1a) as observed in other studies ^50^.

**Figure 1.**
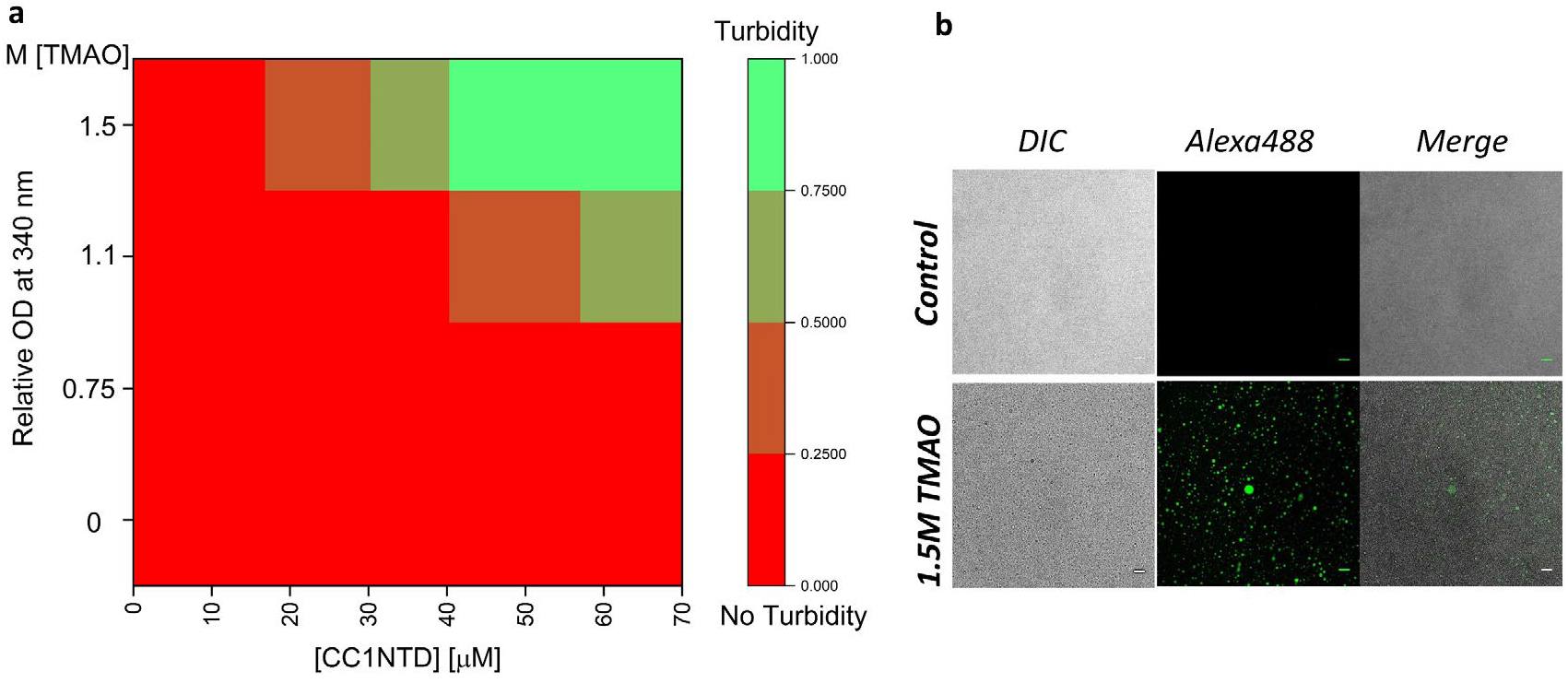
CC1NTD turbidity in the presence of TMAO. (a) UV-visible spectrophotometry of the concentration dependence of CC1NTD turbidity at different TMAO concentrations (0, 0.75, 1.1, and 1.5M) added to buffer (50mM Tris-HCl, 150mM NaCl, 2mM TCEP, pH 8.0) measured at 340nm. Values are normalized relative to the highest value. (b) DIC confocal images of Alexa Fluor 488-labeled CC1NTD (2.6µM (4% molar ratio) in 60µM of unlabeled CC1NTD) without (top) and with 1.5M TMAO showing that CC1NTD condenses to form spherical condensates in the presence of TMAO. Scale bar – 10µm

Results from confocal microscopy revealed that these turbid solutions contained spherical condensates. In the presence of 1.5M TMAO, CC1NTD formed droplets with an average diameter of 1.86 ± 0.13µm (Fig. 1b). Confocal imaging of mixtures containing Alexa Fluor 488-labeled CC1NTD and unlabeled CC1NTD showed that fluorescence was strongly enriched within the droplets relative to the surrounding solution, indicating preferential partitioning of CC1NTD into the dense phase. Consistent with this, the apparent partition coefficient was 41.3 ± 3.0, reflecting substantial enrichment of CC1NTD within condensates, and supernatant depletion measurements showed that approximately 40% of the protein was present in the condensed phase.

To assess whether these condensates displayed liquid-like behavior, we monitored them by time-lapse imaging. CC1NTD droplets underwent fusion and changed morphology over time (Movie S1), behavior consistent with dynamic, liquid-like condensates rather than static aggregates. Together, these data show that under reducing conditions CC1NTD forms TMAO-dependent, protein-enriched condensates with liquid-like properties.

### TMAO compacts CC1NTD conformation without folding it

Small-angle X-ray scattering (SAXS) was used to examine the conformation of CC1NTD in the presence of TMAO. At 0.75M TMAO, 60µM CC1NTD did not undergo LLPS, and the SAXS profile showed no evidence of higher-order assembly in the low-*Q* region. At higher TMAO concentrations (1.1 and 1.5M), where LLPS was observed by DIC microscopy, a Porod-like upturn appeared at low *Q*, consistent with the emergence of larger structures ^51^. From Guinier analysis, (Supplementary Information S6, Fig. S2) the radius of gyration (*R*_g_) of CC1NTD decreased from 29.7 ± 0.4 Å without TMAO, to 24.9 ± 0.3 Å at 0.75M TMAO, and 23.5 ± 0.6 Å at 1.1M TMAO (Fig. 2a; Table 1), indicating that CC1NTD undergoes compaction at sub-LLPS concentrations of TMAO. However, interparticle interference effects between CC1 chains may contribute to a smaller *R_g_* values at higher TMAO concentrations. Despite this compaction, Kratky analysis showed no major change in overall conformation, indicating that CC1NTD remains disordered (Fig. 2b). Circular dichroism measurements supported this conclusion, supporting that CC1NTD does not adopt stable secondary structure in the presence of TMAO (Fig. 2c).

**Figure 2.**
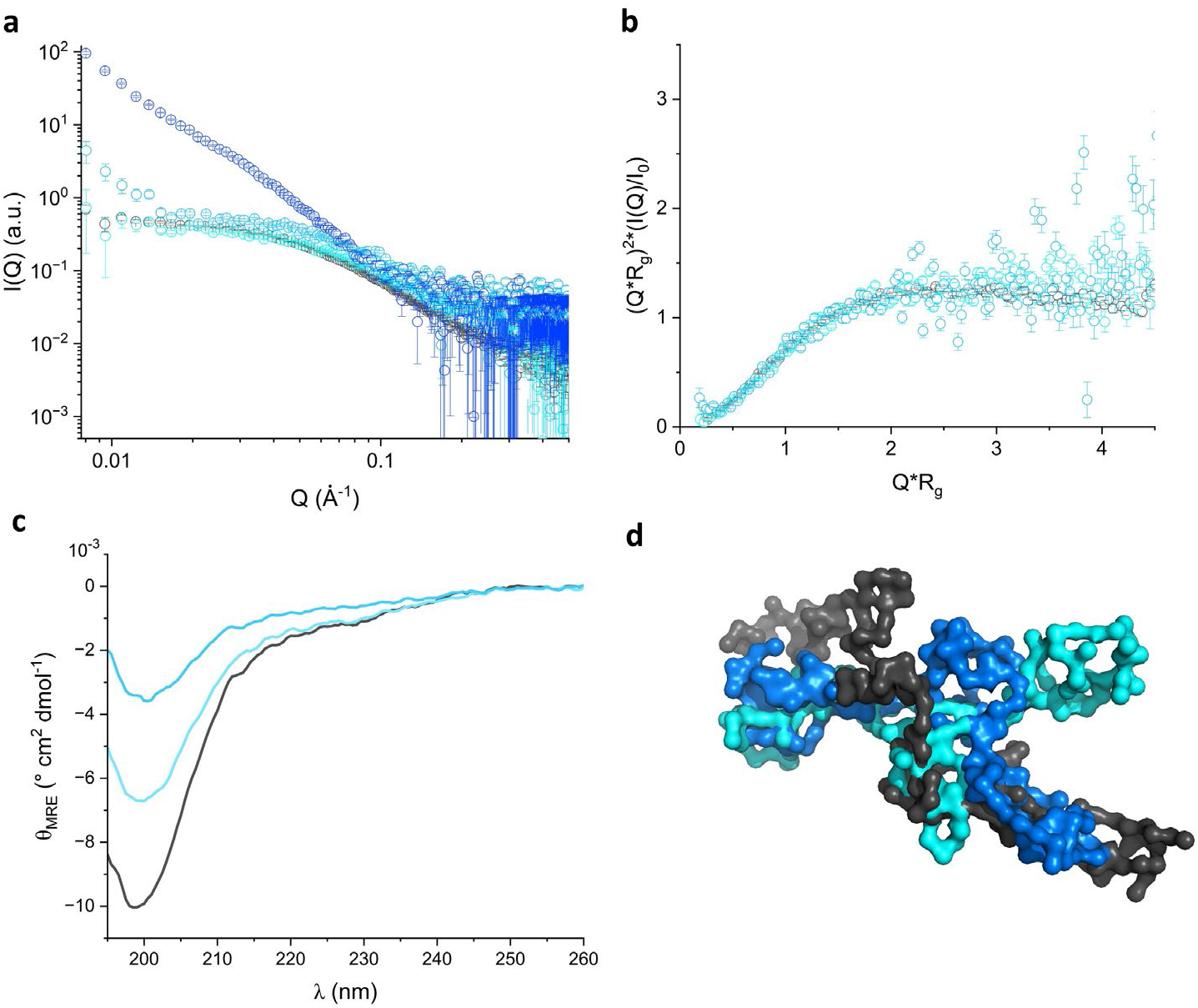
SAXS analysis of CC1NTD in the presence of TMAO (a) SAXS of CC1NTD (60µM) in the presence of 0M (open black circles), 0.75M (open cyan circles), 1.1M (open light blue circles), and 1.5M (open dark blue circles) TMAO; (b) Kratky representations of CC1NTD in TMAO concentrations of 0M (open black circles), 0.75M (open cyan circles), 1.1M (open light blue circles) showing that CC1NTD retains its unfolded conformation; (c) Circular dichroism spectroscopy of CC1NTD in 0M (black), 0.75M (cyan), and 1.1M (light blue) TMAO; (d) GASBOR ab initio reconstruction of CC1NTD generated from SAXS data in 0M (black), 0.75M (cyan), and 1.1M (blue) TMAO.

**Table 1.** Structural parameters of CC1NTD condensates obtained from SAXS.

|  | Model-independent |  | Model-dependent |  |  |  |  |
| --- | --- | --- | --- | --- | --- | --- | --- |
| [TMAO] (M) | Guinier | Real space | Debye |  |  | Excluded volume |  |
| | $R_g$ (Å) | $R_g$ (Å) | $R_g$ (Å) | Persistence length (Å) | Amino acids per length | $R_g$ (Å) | Porod exponent |
| 0 | $29.7 \pm 0.4$ | $30.8 \pm 0.2$ | $33.5 \pm 0.1$ | 8.3 | 2.2 | $32.8 \pm 0.2$ | 2.2 |
| 0.75 | $24.9 \pm 0.3$ | $25.6 \pm 0.2$ | $26.8 \pm 0.4$ | 4.9 | 1.6 | $29.9 \pm 0.2$ | 1.9 |
| 1.1 | $23.5 \pm 0.6$ | $24.9 \pm 0.1$ | $26.6 \pm 0.7$ | 4.7 | 1.3 | $29.3 \pm 0.4$ | 1.8 |

Guinier analysis alone is insufficient to describe the conformational heterogeneity of intrinsically disordered proteins, we also analyzed the data using the pair-distance distribution function, the Debye Gaussian coil model (Eq. 1), and a polymer excluded-volume model. All three approaches supported TMAO-induced compaction (Table 1). Real-space *R*_g_ values decreased by about 17%, and Debye model fits showed a similar decrease in *R*_g_, together with a reduction in persistence length and in the number of amino acids per persistence length (Eq. 2), consistent with chain compaction. Fits to the excluded-volume model also showed a modest decrease in *R_g_*, while Porod exponents between 1.8 and 2.2 indicated that CC1NTD remained disordered across all conditions and transitions from an ideal random coil to a large network of random coils (or swollen Gaussian coil) ^52,53^. GASBOR reconstructions likewise showed a more compact envelope in the presence of TMAO ^40^. Together, these results indicate that TMAO compacts CC1NTD before and during condensate formation but does not induce folding.

### CC1NTD condensates interact with tubulin and accelerate microtubule polymerization

CC1NTD has previously been shown to bundle microtubules ^21,22^, prompting us to test whether CC1NTD condensates remain functionally active toward tubulin. Alexa Fluor 488-labeled CC1NTD condensates were formed in 1.5M TMAO and then mixed with HiLyte 647-labeled tubulin. Confocal microscopy showed that tubulin partitioned into CC1NTD condensates (Fig. 3a). Upon tubulin addition, the average condensate diameter increased from 1.5 to ∼ 3µm, consistent with tubulin recruitment into the dense phase (Fig. 3b).

**Figure 3.**
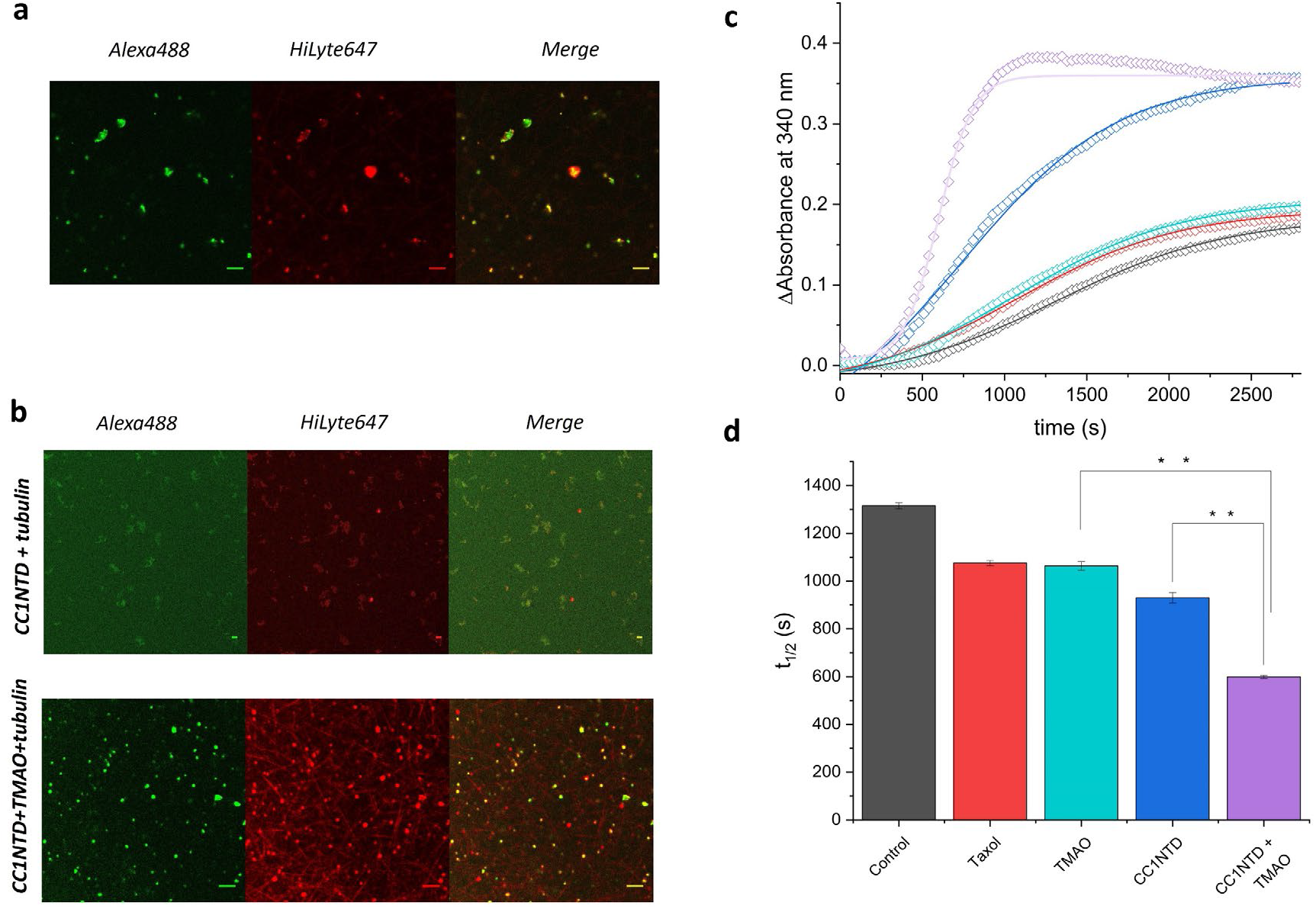
Microtubule polymerization by CC1NTD condensates (a) Confocal microscopy of CC1NTD and tubulin. The 488nm fluorescence, 647nm fluorescence, and merged images of Alexa Fluor 488-labeled CC1NTD and HiLyte 647 tubulin in presence of 1.5M TMAO showing tubulin partitioning with CC1NTD. Scale bar – 10 µm; (b) 488nm fluorescence, 647nm fluorescence, and merged images of Alexa Fluor 488 labeled CC1NTD and HiLyte 647 tubulin collected after 10min in the presence of polymerization buffer under different conditions: (top) Alexa Fluor 488-labeled CC1NTD in osmolyte-free conditions colocalizing with HiLyte 647 tubulin with no evidence of microtubules formation; (bottom) Alexa Fluor 488-labeled CC1NTD condensates in 1.5M TMAO showing rod-shaped microtubules. Scale – 10µm; (c) Polymerization assay of microtubules shown as turbidity measured at 340nm vs. time (s) under different conditions. Microtubule polymerization was performed in the presence of CC1NTD condensates and 1.5M TMAO (open violet diamonds) and CC1NTD without TMAO (open blue diamonds). The control measurements for microtubule polymerization were performed in the absence of CC1NTD with 10µM paclitaxel (open cyan diamonds), 1.5M TMAO (open red diamonds), and general tubulin buffer (open black diamonds). n = 2 experiments. Data shown as mean ± sem. The solid lines are the exponential Boltzmann fits; (d) Bar chart comparing t_1/2_ values obtained from microtubule polymerization assay fits. ** unpaired t-test p-value < 0.0001.

This was followed by investigating how CC1NTD-tubulin interactions affected microtubule assembly. In turbidity-based polymerization assays, tubulin polymerized substantially faster in the presence of CC1NTD condensates formed with 1.5 M TMAO compared to CC1NTD alone (Figs. 3c,d; Table 2). The polymerization curves were fitted with a Boltzmann sigmoidal function (Eq. 3) to obtain the polymerization half-time 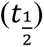 and τ, the characteristic time scale that describes thesharpness of the polymerization transition, with smaller values corresponding to a more abrupt increase in polymerization signal. The polymerization half-times were 598.8 ± 6.4s for CC1NTD condensates in 1.5M TMAO, compared with 929.8 ± 21s for CC1NTD without TMAO and 1063-1315s for the control conditions (Table 2). The corresponding τ values showed a lower value for CC1NTD condensates, ∼ 105s, indicating a steeper and more abrupt increase in turbidity during microtubule assembly. Consistent with these measurements, confocal imaging after 10min showed rod-shaped microtubules in the presence of CC1NTD condensates, whereas no visible microtubules were observed with CC1NTD under osmolyte-free conditions (Fig. 3b). Together, these data indicate that CC1NTD condensates accelerate microtubule polymerization.

**Table 2.** The effect of Microtubule polymerization assay parameters.

| Conditions | Half-time ($t_{1/2}$) (s) | $\tau$ (s) | Total polymer mass (O.D.) |
| --- | --- | --- | --- |
| Buffer only | $1315.3 \pm 12.4$ | $519.1 \pm 11.4$ | $0.17 \pm 0.09$ |
| 10 $\mu\text{M}$ Paclitaxel | $1074.8 \pm 10.5$ | $499.3 \pm 8$ | $0.18 \pm 0.01$ |
| 1.5 M TMAO | $1063 \pm 18$ | $510.5 \pm 13.6$ | $0.19 \pm 0.04$ |
| CC1NTD alone | $929.8 \pm 21$ | $312.2 \pm 19.3$ | $0.35 \pm 0.02$ |
| CC1NTD + 1.5 M TMAO | $598.8 \pm 6.4$ | $104.8 \pm 5.3$ | $0.35 \pm 0.07$ |

To examine how tubulin affects condensate behavior, we characterized CC1NTD condensates with and without tubulin using time-resolved dynamic light scattering (trDLS) ^54^ and FRAP measurements (Fig. 4). In 1.5M TMAO, CC1NTD alone formed spherical particles with a hydrodynamic radius of approximately 0.9µm, consistent with the microscopy measurements (0.93 µm radius). To quantify condensate growth, the trDLS data were analyzed using a power law (Eq. S3, S4) derived from the Ostwald ripening relationship (*R*_h_ ∝ *t*^b^, see Supplementary Information) ^55–57^. The fitted coarsening exponent for CC1NTD condensates (*b* = 0.17 ± 0.01) was lower than the classical value for diffusion-limited ripening, consistent with relatively slow condensate growth ^54^. In the presence of tubulin, the hydrodynamic radius increased transiently to about 1.4 µm before declining after ∼600s (Fig. 4a), a timescale that closely matched the polymerization half-time measured in the turbidity assay. The observed decline may be because the microtubules size became too large to be captured by DLS or that the particles settled out of solution. FRAP measurements showed that CC1NTD within condensates recovered with a half-time of 25.6 ± 1.2s (Eq. S5) and an apparent diffusion coefficient of 0.021 ± 0.003µm^2^/s (Eq. S6). In contrast, CC1NTD in tubulin-associated condensates recovered much more rapidly (Fig. 4b), with a half-time of 4.5 ± 0.1s and an apparent diffusion coefficient of 0.08 ± 0.001µm^2^/s. These results indicate that tubulin association increases the internal dynamics of CC1NTD condensates. Overall, microscopy, polymerization, trDLS, and FRAP data support a consistent model in which CC1NTD condensates recruit tubulin, promote more rapid microtubule assembly, and adopt a more dynamic state during tubulin interaction.

**Figure 4.**
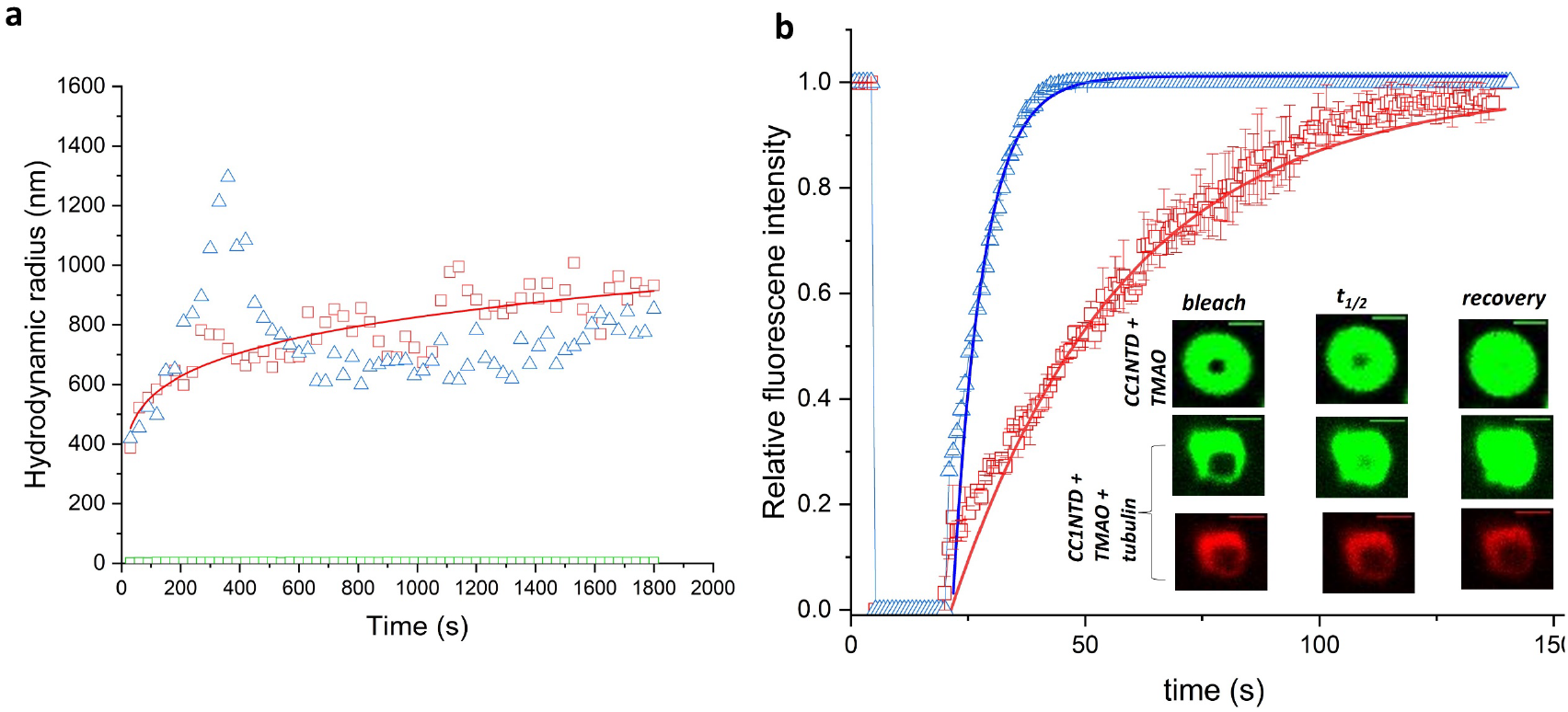
Characterization of CC1NTD condensate dynamics (a) trDLS of hydrodynamic radius evolution of CC1NTD condensates over time. CC1NTD condensates (1.5M TMAO) with the power law fitting (red line); CC1NTD condensates with tubulin in polymerization buffer (open blue triangles); and CC1NTD in the absence of TMAO remains unchanged in size (open green squares); (b) FRAP assay of CC1NTD condensates in 1.5M TMAO with and without tubulin. Alexa Fluor 488-labeled CC1NTD condensates shown as open red squares fit to an exponential function (red line). Alexa Fluor 488-CC1NTD labeled condensates mixed with HiLyte 647 labeled tubulin in polymerization buffer (open blue triangles). The exponential fit is shown as a blue line. Inset (top) 488 fluorescence channel images of Alexa Fluor 488-labeled CC1NTD condensates during bleaching, at half-time recovery, and after recovery at 140s; (middle and bottom) Alexa Fluor 488-CC1NTD condensates in the presence of HiLyte 647 tubulin from 488 (middle) and 647 (bottom) fluorescence channels during bleaching, at half-time recovery, and after recovery at 140s. Scale bar – 2µm; n = 3 experiments.

### Contrast-variation SANS reveals hierarchical organization within CC1NTD–tubulin condensates

Small-angle neutron scattering with contrast variation was used to examine the internal organization of CC1NTD–tubulin condensates. In 85% D_2_O, the scattering contribution from deuterated CC1NTD (dCC1NTD) was minimized, so the measured signal was dominated by protiated tubulin. In 42% D_2_O, the scattering contribution from protiated tubulin was minimized, so the signal predominantly emphasized dCC1NTD. The fully protiated CC1NTD–tubulin sample measured in 85% D_2_O contained scattering contributions from both components. The profiles were described using a combined Unified fit and core–shell cylinder model, in which the Unified term captured mesoscale organization and the cylinder term described smaller elongated structures (See Materials and Methods and Supplementary Information) (Fig. 5).

**Figure 5.**
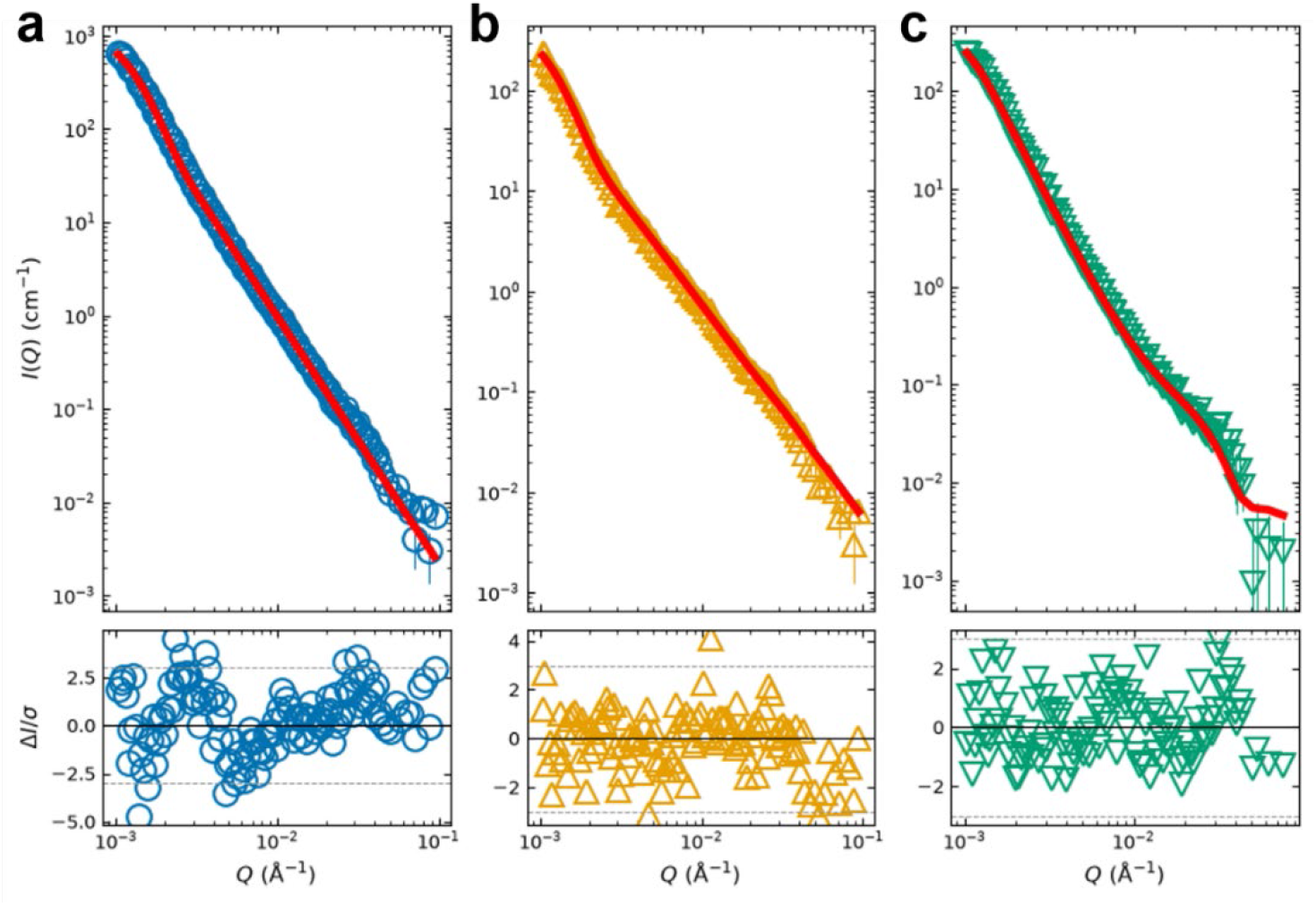
Contrast-variation SANS of CC1NTD–tubulin condensates. **(a)** SANS profiles and corresponding model fits are shown for protiated CC1NTD–tubulin condensates in 85% D₂O (open blue circles); (b) dCC1NTD–tubulin condensates in 85% D₂O (yellow triangles); and dCC1NTD–tubulin condensates in 42% D₂O (green inverted triangles). Error bars indicate the uncertainties from neutron counting statistics. The combined Unified Fit and core– shell cylinder model is shown as a solid red line. Normalized residuals are plotted beneath each profile using the same color scheme as the main plots. Figure S3 shows the profiles for the long and short instrument configuration and the contribution of each structural model to the data.

For the fully protiated CC1NTD–tubulin sample measured in 85% D_2_O, both proteins contributed to the scattering. The Unified fit gave a contrast-weighted *R_g_* of 165 ± 1nm and a power-law exponent *P* = 2.6 ± 0.1 (Fig. 5, Table 3). Because this exponent falls within the mass-fractal range, the data are interpreted as an interconnected, compact mass-fractal-like organization. The *R_g_*-value is interpreted as lower bound or characteristic scale of the condensates rather than a precisely defined domain radius because the Guinier range was outside the limits (*Q_min_* = 0.001Å^-1^). However, the large size does confirm scattering from larger clusters. Contrast variation revealed large-scale organization for the tubulin- and CC1NTD-dominated components. In 85% D_2_O, where the contribution from dCC1NTD was minimized, the tubulin-dominated profile gave an *R_g_* = 180 ± 3nm and *P* = 2.1 ± 0.1 (Fig. 5, Table 3). The corresponding apparent mass-fractal dimension, *D_m_* = 2.1, is consistent with a loose, branched network rather than a densely packed three-dimensional structure. Similar to the interpretation of the full contrast condition the *R_g_* value is interpreted as lower boundary for size of tubulin assemblies. In 42% D_2_O, where the scattering is dominated by dCC1NTD, the fit gave a similar *R_g_* of 175 ± 4nm and *P* of 3.2 ± 0.1 (Fig. 5, Table 3). This exponent is consistent with surface-fractal-like or interface-dominated scattering. The corresponding apparent surface-fractal dimension, (*D_s_* = 6 − *P* = 2.8), is consistent with a rough surface or heterogeneous CC1NTD-rich environment. Similar to the other conditions, the *R_g_* is interpreted as lower bound or characteristic scale of CC1NTD condensates rather than a precisely defined domain radius.

**Table 3.** Fit parameters for CC1NTD–tubulin condensates.

| **Sample and contrast condition** | **Dominant contrast contribution** | **$R_g(\text{nm})$** | **$P$** | **Fractal interpretation** |
| --- | --- | --- | --- | --- |
| Protiated CC1NTD–tubulin in 85% D2O | CC1NTD and tubulin | $165 \pm 1$ | $2.6 \pm 0.1$ | Mass-fractal-like, relatively compact and interconnected; $D_m \approx 2.65$ |
| dCC1NTD–tubulin in 85% D2O | Predominantly tubulin | $180 \pm 3$ | $2.1 \pm 0.1$ | Open, branched mass-fractal-like organization; $D_m \approx 2.11$ |
| dCC1NTD–tubulin in 42% D2O | Predominantly CC1NTD | $175 \pm 4$ | $3.2 \pm 0.1$ | Surface-fractal-like or interface-dominated; $D_s \approx 2.8$ |

At smaller length scales, the data were fitted to an extended core–shell cylinder model and yielded a core radius of 8.4 ± 0.1nm (16.7 ± 0.3nm diam.) and shell thickness of 5.8 ± 0.3nm, giving an overall diameter of 28.4 ± 0.6nm (Fig. 5, Table 3). This diameter is comparable with the microtubule-like cross-section reported previously ^21^. The high-*Q* cylindrical feature was retained under the tubulin-dominated contrast but was reduced or absent under the CC1NTD-dominated contrast, supporting that it is associated with the tubulin-rich component. Together, the contrast-variation data support hierarchical organization in which tubulin-rich cylindrical assemblies participate in an open mesoscale network within a broader and more interface-rich CC1NTD environment (Table 3). This is consistent with a multicomponent condensate containing distinct but overlapping CC1NTD- and tubulin-rich structural regions.

### CC2NTD does not phase separate independently but partitions into CC1NTD condensates

Arabidopsis CC2 is a closely related paralog of CC1, sharing 84% sequence identity in the N-terminal disordered domain and, like CC1NTD, is predicted to be intrinsically disordered (Fig. S4a.b). However, CC2NTD is shorter than CC1NTD and lacks one of the microtubule-interacting regions previously identified in CC1NTD ^22^. CD shows that CC2NTD is predominantly disordered but has increased β-sheet content and a lower amount of disordered or random coil structural features compared to CC1NTD (Supplementary Information Section S8, Fig. S4c, Table S2). SAXS analyses further supported a disordered conformation for CC2NTD in solution, with structural parameters broadly similar to those of CC1NTD (Fig. S4d,e; Table S3). Further functional analyses of CC2NTD clearly showed that it shares the same ability as CC1NTD to interact with microtubules and assemble them into well-ordered arrays (Supplementary Information Sections S8-12, Fig. S5; Table S4). Together, these data establish the suitability of CC2NTD for testing which features of CC1NTD are specifically associated with condensate formation.

We investigated whether CC2NTD undergoes phase separation under the same conditions that promote CC1NTD condensation. Although the same in silico analyses used for CC1NTD suggested that CC2NTD has phase-separation propensity, CC2NTD showed no detectable turbidity or condensate formation in 1.5M TMAO under the conditions used for CC1NTD (Fig. 6a-c). This contrasts with CC1NTD, which readily formed condensates under the same conditions, indicating that high sequence similarity and intrinsic disorder alone are not sufficient to drive condensation in this system. CC2NTD retained other biophysical features of CC1NTD, supporting the conclusion that its major difference under these conditions lies in condensate formation rather than global disorder.

**Figure 6.**
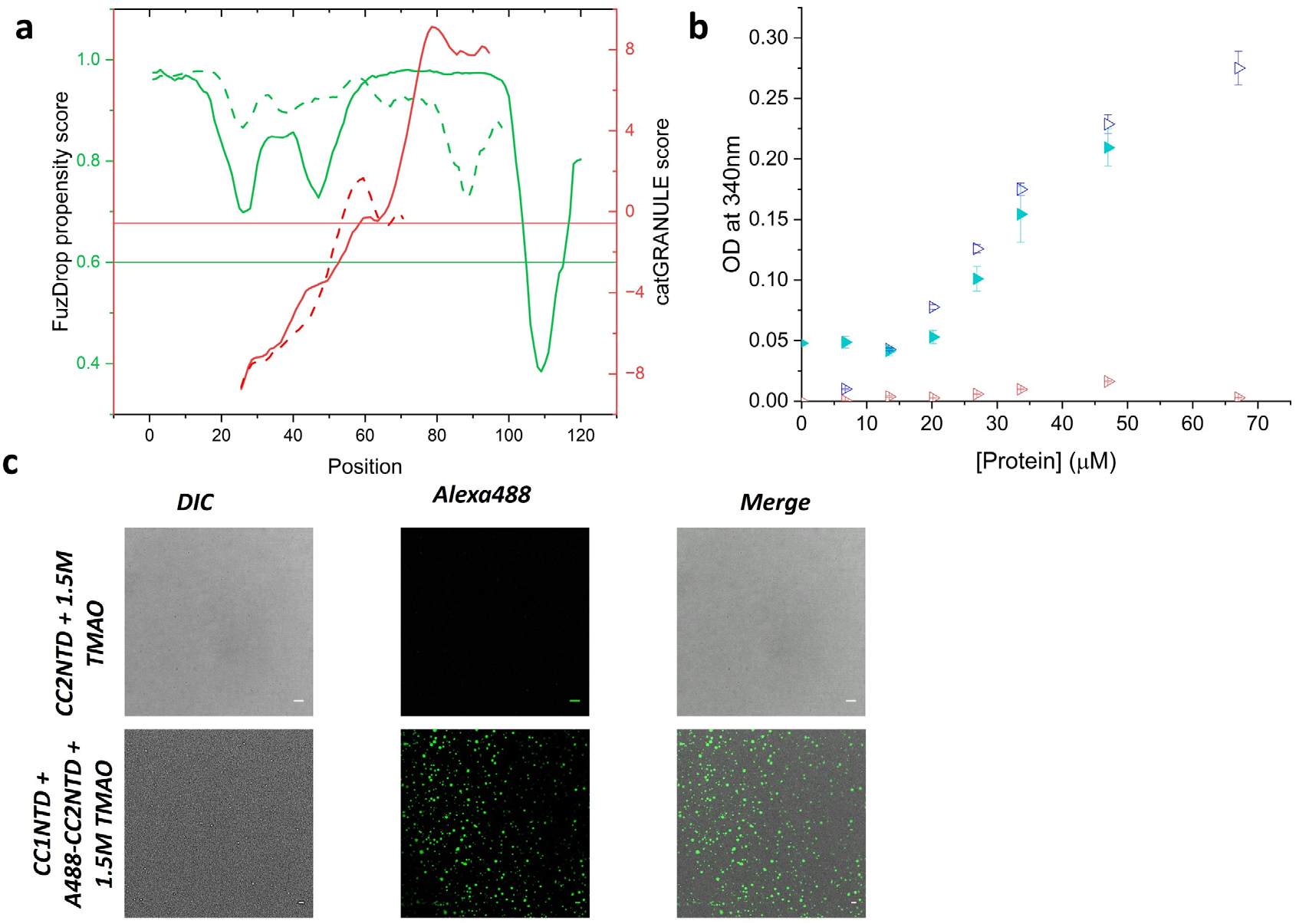
Characterization of CC2NTD propensity to form LLPS and its interactions with CC1NTD condensates (a) Phase separation propensity of CC2NTD and comparison with CC1NTD. The plot shows results from FuzDrop algorithm for CC1NTD (green) and CC2NTD (dashed green) and catGRANULE algorithm for CC1NTD (red) and CC2NTD (dashed red). Horizontal green and red lines indicate the cutoff above is the positive prediction for LLPS propensity; (b) Turbidity measurements showing that CC1NTD (open blue triangles) becomes turbid whereas CC2NTD (open red triangles) does not. However, addition of CC1NTD to 60µM CC2NTD in 1.5M TMAO (closed cyan triangles) results in a turbid solution; (c) Confocal imaging of CC2NTD in 1.5M TMAO of (top) Alexa Fluor 488-labeled CC2NTD in 1.5M TMAO and (bottom) Alexa Fluor 488-labeled CC2NTD localizing into CC1NTD condensates in the presence of 1.5M TMAO. Scale bar – 10 µm.

Although CC2NTD did not phase-separate on its own, it did partition into CC1NTD condensates. Adding increasing amounts of CC1NTD (up to 60µM) to a fixed concentration of CC2NTD (60µM) in 1.5M TMAO, resulted in a turbid solution as shown in Fig. 6b. When Alexa Fluor 488-labeled CC2NTD was mixed with unlabeled CC1NTD in 1.5M TMAO, confocal microscopy showed clear enrichment of CC2NTD within CC1NTD condensates (Fig. 6c). These mixed condensates had an average diameter of 1.5µm, similar to that of CC1NTD condensates alone and the apparent partition coefficient of CC2NTD was 164.5 ± 28.3, indicating strong recruitment into the dense phase. The mixed condensates also remained dynamic, exhibiting fusion and time-dependent morphological changes similar to those observed for CC1NTD alone (Movie S2). This behavior is consistent with the possibility that CC2NTD can be recruited into a CC1NTDdense phase despite lacking independent condensate-forming activity under the conditions tested. Together, these results identify CC2NTD as a non-condensing, but condensate-partitioning in the presence of CC1NTD, under the conditions tested.

## Discussion

In this study, we show that the intrinsically disordered N-terminal domain of CC1 forms condensates in the presence of TMAO, a natural osmolyte associated with plant stress responses. Condensate formation was both protein and TMAO-dependent. Higher TMAO concentrations lowered the CC1NTD concentration threshold for turbidity and droplet formation. Microscopy, fluorescence partitioning, and supernatant depletion further showed that these condensates are enriched in CC1NTD, while time-lapse imaging demonstrated fusion and morphology changes consistent with liquid-like behavior. These observations support the interpretation that CC1NTD undergoes liquid-like phase separation under the tested conditions rather than forming static aggregates. Prior osmolyte studies indicate that in vitro measurements of osmolyte effects require the osmolyte concentration to remain high relative to the protein concentration, consistent with preferential exclusion models ^58,59^. In that framework, the inverse relationship observed here between CC1NTD and TMAO concentrations is consistent with crowding or osmolyte-assisted condensation rather than a discrete ligand-binding effect. Thus, the present data support the conclusion that TMAO can shift the conformational and assembly landscape of CC1NTD toward condensate formation under reducing conditions.

A second important result is that TMAO promotes compaction of CC1NTD without inducing folding. SAXS showed a decrease in overall size at sub-LLPS and LLPS-promoting TMAO concentrations, and model-based analysis further supported chain compaction. However, Kratky analysis, Porod exponents, and CD all indicated that CC1NTD remained disordered across these conditions. These findings suggest that phase separation is accompanied by remodeling of the disordered ensemble rather than by acquisition of stable secondary or tertiary structure. In other words, CC1NTD does not appear to condense because it folds; instead, it remains intrinsically disordered while sampling a more compact conformational state that favors condensation.

Another insight from this work is that CC1NTD condensates remain functionally active toward tubulin. Tubulin partitioned into CC1NTD condensates, and this interaction was followed by faster microtubule assembly in turbidity-based polymerization assays. The clearest functional metric was the reduction in polymerization half-time in the presence of CC1NTD condensates relative to CC1NTD alone and to control conditions. The data therefore support the view that condensate formation may be coupled to tubulin-related function. Under in vitro salt-stress-mimicking conditions, increased local concentration of CC1NTD within condensates may enhance tubulin engagement and accelerate microtubule formation. The confocal imaging results are also consistent with this interpretation. Time-resolved DLS showed that CC1NTD condensates undergo relatively slow growth on their own but exhibit a transient increase in hydrodynamic size upon tubulin addition, on a timescale similar to the measured polymerization half-time. FRAP further showed that CC1NTD recovers more rapidly in tubulin-associated condensates than in CC1NTD-only condensates. This behavior is consistent with a more dynamic condensate state in the presence of tubulin. One reasonable interpretation is that condensation increases the effective local concentration of CC1NTD and creates a multivalent environment that favors the recruitment of tubulin and rapid microtubule formation. This interpretation is also consistent with previous evidence that CC1NTD engages tubulin or microtubules through multiple interaction sites ^22^. Contrast-variation SANS results support formation of a multicomponent CC1NTD–tubulin condensate rather than simple adsorption of tubulin onto the exterior of a CC1NTD droplet. Within this mixed phase, the two components appear to adopt distinct but overlapping spatial organizations, with tubulin-rich cylindrical assemblies embedded in a broader CC1-rich environment. Similar coupling between condensate formation and tubulin-dependent assembly has been described for other intrinsically disordered microtubule-associated systems, including tau, suggesting that condensation may represent a general mechanism for locally enhancing microtubule assembly reactions ^54,60^.

The behavior of CC2NTD is informative for understanding what is specific to CC1NTD condensation. CC2NTD is highly similar to CC1NTD in sequence and, like CC1NTD, remains disordered in solution by CD and SAXS. Nevertheless, CC2NTD did not show detectable turbidity or condensate formation under the same TMAO-containing conditions that readily induced CC1NTD condensation. This indicates that intrinsic disorder and broad sequence similarity are not sufficient to drive condensation in this system. Instead, condensate formation appears to depend on more specific sequence features. Several sequence-level differences may contribute to this difference. First, a tyrosine residue in CC1NTD is replaced by phenylalanine in CC2NTD, which could alter the aromatic interactions that often contribute to phase separation ^61–63^. Second, CC2NTD is shorter overall, which may reduce the effective valency or spatial distribution of weak interactions ^62,64,65^. Third, CC2 lacks a region present in CC1 spanning approximately residues 62-85/86 that is enriched in charged residues. Its absence may reduce the electrostatic and multivalent interactions needed to support phase separation under the tested conditions. These ideas remain hypotheses rather than demonstrated mechanisms, but together they provide a plausible framework for why CC2NTD fails to condense independently despite its overall similarity to CC1NTD. At the same time, CC2NTD was strongly recruited into CC1NTD condensates. This result is important because it shows that a closely related non-condensing protein can still partition into a CC1NTD-defined dense phase. This suggests that CC1NTD creates a condensate environment that can recruit CC2NTD through shared interaction features, even though CC2NTD does not independently satisfy the requirements for phase separation under these conditions. This behavior is consistent with, but does not prove, a scaffold-client type relationship between the two proteins. A recent study proposed a similar scaffold-client mechanism for the phase-separation behavior of the heterochromatin HP1α, HP1β, and HP1γ paralogs ^66^.

### Biological implications for CC condensates in plants

The broader significance of these findings lies in providing a physicochemical framework for how CC proteins function during stress. Previous work has implicated CC1 in sustaining cellulose synthesis under salt stress and has shown that CC1 accumulates in small CESA compartments (smaCCs) or microtubule-associated cellulose synthase compartments (MASCs), which are stress-associated cellulose synthase compartments ^18^. The present data do not demonstrate that smaCCs/MASCs are LLPS-derived structures, but they do raise the possibility that stress-responsive condensation of the CC1 N-terminal domain could contribute to compartmentalization, microtubule interactions, or local assembly reactions in ways relevant to these compartments. In this scenario, condensation of CC1 could help concentrate CC1 locally, recruit tubulin, and promote a more efficient microtubule recovery response under stress conditions. Because CC1 is linked to cellulose synthase stress biology, such a mechanism could in principle aid the stabilization or organization of cellulose synthase-associated compartments during stress adaptation. This model is also compatible with the idea that the disordered cytosolic regions of CC1 and possibly other cellulose synthase-associated proteins contribute to transient compartment formation or stabilization ^67^. Future studies of full-length CC proteins and other CSC proteins with similar phase separation properties and investigating roles of posttranslational modifications of these proteins on their phase behavior may pave way for a better understanding of CSC compartmentalization ^68^.

Several important questions remain unresolved. It is not yet known whether CC1 undergoes comparable condensation in vivo, whether TMAO is itself the relevant physiological trigger or instead at very high concentrations serves as a useful experimental proxy for a broader stress-induced physicochemical state, or whether full-length CC1 behaves similarly to the isolated N-terminal domain. It also remains unclear whether CESA proteins or other cellulose synthase-associated factors co-condense with CC1, and whether condensation directly contributes to smaCC/MASC formation, stabilization, or recovery. Addressing these questions will be necessary to connect the present in vitro biophysical model more directly to plant stress physiology.

## Conclusions

CC1NTD forms protein-enriched, liquid-like condensates under reducing, TMAO-containing conditions and undergoes compaction without folding as it enters this condensed state. These condensates recruit tubulin, remain dynamic, and accelerate microtubule polymerization, linking condensate formation to a measurable functional consequence. Comparison with CC2NTD further shows that close sequence similarity and intrinsic disorder are not sufficient for condensation, while also indicating that related proteins can still be recruited into a CC1NTD-defined dense phase. Together, these findings support a model in which stress-responsive condensation of CC1 provides a plausible physicochemical framework for understanding how CC proteins could contribute to microtubule regulation and cellulose synthase-associated stress responses.

## Supporting information

additional methods and results

## Supporting Information

Additional details on protein expression, purification and labeling; sample preparation for microscopy; dynamic light scattering methods; small-angle X-ray scattering Guinier analysis; small-angle neutron scattering data fitting approach; and CC2NTD characterization including bioinformatic and solution studies, microtubule binding studies; and time lapse movies of CC1NTD condensates and CC1NTD/CC2NTD co-condensates.

## Acknowledgements

VG and HO’N acknowledge funding by Center for Lignocellulose Structure and Formation supported by the U.S. Department of Energy (DOE), Office of Science, Basic Energy Sciences under Award # DE-SC0001090. SANS studies of CC1NTD LLPS were supported by the Laboratory Directed Research and Development Program of Oak Ridge National Laboratory under IPTS #35830. Bio-SANS is part of the Center for Structural Molecular Biology funded by DOE Office of DOE Biological & Environmental Research (OBER) project ERKP291. RL acknowledges support from the National Institutes of Health R35GM142946. This research used resources at the High Flux Isotope Reactor and Spallation Neutron Source, a DOE Office of Science User Facility operated by the Oak Ridge National Laboratory.

## Author contributions

VG designed and performed the experiments, and associated data analysis. WCL, ACH, QZ planned and performed SANS experiments and data analysis. RL and JLK performed fluorescence and electron microscopy experiments. HO’N directed the research project. VG and HO’N wrote the manuscript with input from all authors. All authors had the opportunity to read and comment on the manuscript.

## Table of Contents Figure

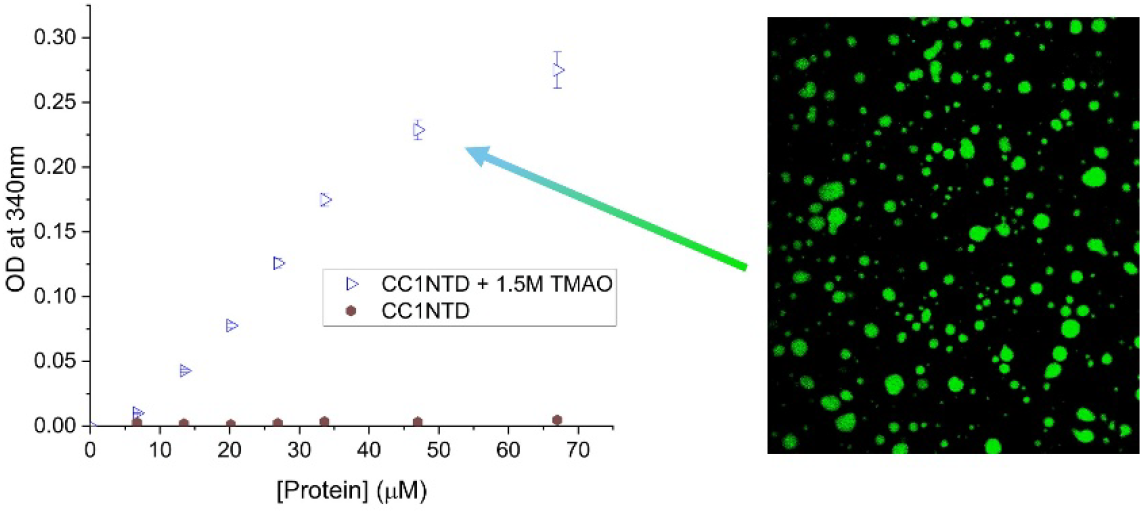

## Notes

### Competing Interest Statement

The authors have declared no competing interest.

## References

(1) Shin, Y.; Brangwynne, C. P. Liquid Phase Condensation in Cell Physiology and Disease. Science 2017, 357 (6357), eaaf4382. 10.1126/science.aaf4382

(2) Alberti, S.; Gladfelter, A.; Mittag, T. Considerations and Challenges in Studying Liquid-Liquid Phase Separation and Biomolecular Condensates. Cell 2019, 176 (3), 419–434. 10.1016/j.cell.2018.12.035

(3) Dragwidge, J. M.; Van Damme, D. Protein Phase Separation in Plant Membrane Biology: More than Just a Compartmentalization Strategy. Plant Cell 2023, 35 (9), 3162–3172. 10.1093/plcell/koad177

(4) Fang, X.; Wang, L.; Ishikawa, R.; Li, Y.; Fiedler, M.; Liu, F.; Calder, G.; Rowan, B.; Weigel, D.; Li, P.; Dean, C. Arabidopsis FLL2 Promotes Liquid–Liquid Phase Separation of Polyadenylation Complexes. Nature 2019, 569 (7755), 265–269. 10.1038/s41586-019-1165-8

(5) Zhu, P.; Lister, C.; Dean, C. Cold-Induced Arabidopsis FRIGIDA Nuclear Condensates for FLC Repression. Nature 2021, 599 (7886), 657–661. 10.1038/s41586-021-04062-5

(6) Huang, S.; Zhu, S.; Kumar, P.; MacMicking, J. D. A Phase-Separated Nuclear GBPL Circuit Controls Immunity in Plants. Nature 2021, 594 (7863), 424–429. 10.1038/s41586-021-03572-6

(7) Zavaliev, R.; Mohan, R.; Chen, T.; Dong, X. Formation of NPR1 Condensates Promotes Cell Survival during the Plant Immune Response. Cell 2020, 182 (5), 1093–1108.e18. 10.1016/j.cell.2020.07.016

(8) Powers, S. K.; Holehouse, A. S.; Korasick, D. A.; Schreiber, K. H.; Clark, N. M.; Jing, H.; Emenecker, R.; Han, S.; Tycksen, E.; Hwang, I.; Sozzani, R.; Jez, J. M.; Pappu, R. V.; Strader, L. C. Nucleo-Cytoplasmic Partitioning of ARF Proteins Controls Auxin Responses in Arabidopsis Thaliana. Mol. Cell 2019, 76 (1), 177–190.e5. 10.1016/j.molcel.2019.06.044

(9) Gutierrez-Beltran, E.; Elander, P. H.; Dalman, K.; Dayhoff, G. W.; Moschou, P. N.; Uversky, V. N.; Crespo, J. L.; Bozhkov, P. V. Tudor Staphylococcal Nuclease Is a Docking Platform for Stress Granule Components and Is Essential for SnRK1 Activation in Arabidopsis. EMBO J. 2021, 40 (17), EMBJ2020105043. 10.15252/embj.2020105043

(10) Zhu, S.; Gu, J.; Yao, J.; Li, Y.; Zhang, Z.; Xia, W.; Wang, Z.; Gui, X.; Li, L.; Li, D.; Zhang, H.; Liu, C. Liquid-Liquid Phase Separation of RBGD2/4 Is Required for Heat Stress Resistance in Arabidopsis. Dev. Cell 2022, 57 (5), 583–597.e6. 10.1016/j.devcel.2022.02.005

(11) Zhang, Y.; Li, Z.; Chen, N.; Huang, Y.; Huang, S. Phase Separation of Arabidopsis EMB1579 Controls Transcription, mRNA Splicing, and Development. PLOS Biol. 2020, 18 (7), e3000782. 10.1371/journal.pbio.3000782

(12) Dorone, Y.; Boeynaems, S.; Flores, E.; Jin, B.; Hateley, S.; Bossi, F.; Lazarus, E.; Pennington, J. G.; Michiels, E.; De Decker, M.; Vints, K.; Baatsen, P.; Bassel, G. W.; Otegui, M. S.; Holehouse, A. S.; Exposito-Alonso, M.; Sukenik, S.; Gitler, A. D.; Rhee, S. Y. A Prion-like Protein Regulator of Seed Germination Undergoes Hydration-Dependent Phase Separation. Cell 2021, 184 (16), 4284–4298.e27. 10.1016/j.cell.2021.06.009

(13) Arif, Y.; Singh, P.; Siddiqui, H.; Bajguz, A.; Hayat, S. Salinity Induced Physiological and Biochemical Changes in Plants: An Omic Approach towards Salt Stress Tolerance. Plant Physiol. Biochem. 2020, 156, 64–77. 10.1016/j.plaphy.2020.08.042

(14) Muchate, N. S.; Nikalje, G. C.; Rajurkar, N. S.; Suprasanna, P.; Nikam, T. D. Plant Salt Stress: Adaptive Responses, Tolerance Mechanism and Bioengineering for Salt Tolerance. Bot. Rev. 2016, 82 (4), 371–406. 10.1007/s12229-016-9173-y

(15) van Zelm, E.; Zhang, Y.; Testerink, C. Salt Tolerance Mechanisms of Plants. Annu. Rev. Plant Biol. 2020, 71, 403–433. 10.1146/annurev-arplant-050718-100005

(16) Zhao, S.; Zhang, Q.; Liu, M.; Zhou, H.; Ma, C.; Wang, P. Regulation of Plant Responses to Salt Stress. Int. J. Mol. Sci. 2021, 22 (9), 4609. 10.3390/ijms22094609

(17) Colin, L.; Ruhnow, F.; Zhu, J.-K.; Zhao, C.; Zhao, Y.; Persson, S. The Cell Biology of Primary Cell Walls during Salt Stress. Plant Cell 2023, 35 (1), 201–217. 10.1093/plcell/koac292

(18) Endler, A.; Kesten, C.; Schneider, R.; Zhang, Y.; Ivakov, A.; Froehlich, A.; Funke, N.; Persson, S. A Mechanism for Sustained Cellulose Synthesis during Salt Stress. Cell 2015, 162 (6), 1353–1364. 10.1016/j.cell.2015.08.028

(19) Ma, J.; Pazos, I. M.; Gai, F. Microscopic Insights into the Protein-Stabilizing Effect of Trimethylamine N-Oxide (TMAO). Proc. Natl. Acad. Sci. 2014, 111 (23), 8476–8481. 10.1073/pnas.1403224111

(20) Wang, C.; Zhang, L.-J.; Huang, R.-D. Cytoskeleton and Plant Salt Stress Tolerance. Plant Signal. Behav. 2011, 6 (1), 29–31. 10.4161/psb.6.1.14202

(21) Gurumoorthy, V.; Hicks, A.; Tiruvadi-Krishnan, S.; Leite, W. C.; Chodankar, S.; Kolape, J. L.; Smith, J. C.; Lamichhane, R.; O’Neill, H. The N-Terminal Domain of COMPANION OF CELLULOSE SYNTHASE1 Promotes Microtubule Array Formation in Arabidopsis. Plant Physiol. 2025, 199 (1), kiaf392. 10.1093/plphys/kiaf392

(22) Kesten, C.; Wallmann, A.; Schneider, R.; McFarlane, H. E.; Diehl, A.; Khan, G. A.; Van Rossum, B.-J.; Lampugnani, E. R.; Szymanski, W. G.; Cremer, N.; Schmieder, P.; Ford, K. L.; Seiter, F.; Heazlewood, J. L.; Sanchez-Rodriguez, C.; Oschkinat, H.; Persson, S. The Companion of Cellulose Synthase 1 Confers Salt Tolerance through a Tau-like Mechanism in Plants. Nat. Commun. 2019, 10 (1), 857. 10.1038/s41467-019-08780-3

(23) Singh, P.; Choudhary, K. K.; Chaudhary, N.; Gupta, S.; Sahu, M.; Tejaswini, B.; Sarkar, S. Salt Stress Resilience in Plants Mediated through Osmolyte Accumulation and Its Crosstalk Mechanism with Phytohormones. Front. Plant Sci. 2022, 13, 1006617. 10.3389/fpls.2022.1006617

(24) Catalá, R.; López-Cobollo, R.; Berbís, M. Á.; Jiménez-Barbero, J.; Salinas, J. Trimethylamine *N* -Oxide Is a New Plant Molecule That Promotes Abiotic Stress Tolerance. Sci. Adv. 2021, 7 (21), eabd9296. 10.1126/sciadv.abd9296

(25) Bennion, B. J.; Daggett, V. Counteraction of Urea-Induced Protein Denaturation by Trimethylamine *N* -Oxide: A Chemical Chaperone at Atomic Resolution. Proc. Natl. Acad. Sci. 2004, 101 (17), 6433–6438. 10.1073/pnas.0308633101

(26) Canchi, D. R.; García, A. E. Cosolvent Effects on Protein Stability. Annu. Rev. Phys. Chem. 2013, 64 (1), 273–293. 10.1146/annurev-physchem-040412-110156

(27) Ufnal, M.; Zadlo, A.; Ostaszewski, R. TMAO: A Small Molecule of Great Expectations. Nutrition 2015, 31 (11–12), 1317–1323. 10.1016/j.nut.2015.05.006

(28) Zou, Q.; Bennion, B. J.; Daggett, V.; Murphy, K. P. The Molecular Mechanism of Stabilization of Proteins by TMAO and Its Ability to Counteract the Effects of Urea. J. Am. Chem. Soc. 2002, 124 (7), 1192–1202. 10.1021/ja004206b

(29) Madeira, F.; Pearce, M.; Tivey, A. R. N.; Basutkar, P.; Lee, J.; Edbali, O.; Madhusoodanan, N.; Kolesnikov, A.; Lopez, R. Search and Sequence Analysis Tools Services from EMBL-EBI in 2022. Nucleic Acids Res. 2022, 50 (W1), W276–W279. 10.1093/nar/gkac240

(30) Mészáros, B.; Erdős, G.; Dosztányi, Z. IUPred2A: Context-Dependent Prediction of Protein Disorder as a Function of Redox State and Protein Binding. Nucleic Acids Res. 2018, 46 (W1), W329–W337. 10.1093/nar/gky384

(31) Hatos, A.; Tosatto, S. C. E.; Vendruscolo, M.; Fuxreiter, M. FuzDrop on AlphaFold: Visualizing the Sequence-Dependent Propensity of Liquid–Liquid Phase Separation and Aggregation of Proteins. Nucleic Acids Res. 2022, 50 (W1), W337–W344. 10.1093/nar/gkac386

(32) Vendruscolo, M.; Fuxreiter, M. Sequence Determinants of the Aggregation of Proteins Within Condensates Generated by Liquid-Liquid Phase Separation. J. Mol. Biol. 2022, 434 (1), 167201. 10.1016/j.jmb.2021.167201

(33) Bolognesi, B.; Lorenzo Gotor, N.; Dhar, R.; Cirillo, D.; Baldrighi, M.; Tartaglia, G. G.; Lehner, B. A Concentration-Dependent Liquid Phase Separation Can Cause Toxicity upon Increased Protein Expression. Cell Rep. 2016, 16 (1), 222–231. 10.1016/j.celrep.2016.05.076

(34) Chu, X.; Sun, T.; Li, Q.; Xu, Y.; Zhang, Z.; Lai, L.; Pei, J. Prediction of Liquid–Liquid Phase Separating Proteins Using Machine Learning. BMC Bioinformatics 2022, 23 (1), 72. 10.1186/s12859-022-04599-w

(35) Vera, J. C. Measurement of Microgram Quantities of Protein by a Generally Applicable Turbidimetric Procedure. Anal. Biochem. 1988, 174 (1), 187–196. 10.1016/0003-2697(88)90534-9

(36) Schneider, C. A.; Rasband, W. S.; Eliceiri, K. W. NIH Image to ImageJ: 25 Years of Image Analysis. Nat. Methods 2012, 9 (7), 671–675. 10.1038/nmeth.2089

(37) Micsonai, A.; Wien, F.; Bulyáki, É.; Kun, J.; Moussong, É.; Lee, Y.-H.; Goto, Y.; Réfrégiers, M.; Kardos, J. BeStSel: A Web Server for Accurate Protein Secondary Structure Prediction and Fold Recognition from the Circular Dichroism Spectra. Nucleic Acids Res. 2018, 46 (W1), W315–W322. 10.1093/nar/gky497

(38) Hopkins, J. B.; Gillilan, R. E.; Skou, S. *BioXTAS RAW* : Improvements to a Free Open-Source Program for Small-Angle X-Ray Scattering Data Reduction and Analysis. J. Appl. Crystallogr. 2017, 50 (5), 1545–1553. 10.1107/S1600576717011438

(39) Doucet, M.; Cho, J. H.; Alina, G.; Bakker, J.; Bouwman, W.; Butler, P.; Campbell, K.; Gonzales, M.; Heenan, R.; Jackson, A.; Juhas, P.; King, S.; Kienzle, P.; Krzywon, J.; Markvardsen, A.; Nielsen, T.; O’Driscoll, L.; Potrzebowski, W.; Ferraz Leal, R.; Richter, T.; Rozycko, P.; Washington, A. SasView Version 4.1, 2017. 10.5281/ZENODO.438138

(40) Receveur-Brechot, V.; Durand, D. How Random Are Intrinsically Disordered Proteins? A Small Angle Scattering Perspective. Curr. Protein Pept. Sci. 2012, 13 (1), 55–75. 10.2174/138920312799277901

(41) Creighton, T. E. *Proteins: Structures and Molecular Properties*, 2. ed., 8. print.; Freeman: New York, 2006.

(42) Anunciado, D.; Rai, D. K.; Qian, S.; Urban, V.; O’Neill, H. Small-Angle Neutron Scattering Reveals the Assembly of Alpha-Synuclein in Lipid Membranes. Biochim. Biophys. Acta BBA - Proteins Proteomics 2015, 1854 (12), 1881–1889. 10.1016/j.bbapap.2015.08.009

(43) Svergun, D. I.; Petoukhov, M. V.; Koch, M. H. J. Determination of Domain Structure of Proteins from X-Ray Solution Scattering. Biophys. J. 2001, 80 (6), 2946–2953. 10.1016/S0006-3495(01)76260-1

(44) Sahu, S.; Herbst, L.; Quinn, R.; Ross, J. L. Crowder and Surface Effects on Self-Organization of Microtubules. *Phys*. Rev. E 2021, 103 (6), 062408. 10.1103/PhysRevE.103.062408

(45) Nielsen, L.; Khurana, R.; Coats, A.; Frokjaer, S.; Brange, J.; Vyas, S.; Uversky, V. N.; Fink, A. L. Effect of Environmental Factors on the Kinetics of Insulin Fibril Formation: Elucidation of the Molecular Mechanism. Biochemistry 2001, 40 (20), 6036–6046. 10.1021/bi002555c

(46) Schummel, P. H.; Gao, M.; Winter, R. Modulation of the Polymerization Kinetics of α/β-Tubulin by Osmolytes and Macromolecular Crowding. ChemPhysChem 2017, 18 (2), 189– 197. 10.1002/cphc.201601032

(47) Heller, W. T.; Urban, V. S.; Lynn, G. W.; Weiss, K. L.; O’Neill, H. M.; Pingali, S. V.; Qian, S.; Littrell, K. C.; Melnichenko, Y. B.; Buchanan, M. V.; Selby, D. L.; Wignall, G. D.; Butler, P. D.; Myles, D. A. The Bio-SANS Instrument at the High Flux Isotope Reactor of Oak Ridge National Laboratory. J. Appl. Crystallogr. 2014, 47 (4), 1238–1246.

(48) Beaucage, G. Small-Angle Scattering from Polymeric Mass Fractals of Arbitrary Mass-Fractal Dimension. J. Appl. Crystallogr. 1996, 29 (2), 134–146. 10.1107/S0021889895011605

(49) Beaucage, G. Approximations Leading to a Unified Exponential/Power-Law Approach to Small-Angle Scattering. J. Appl. Crystallogr. 1995, 28 (6), 717–728. 10.1107/S0021889895005292

(50) Mondal, S.; Narayan, K.; Botterbusch, S.; Powers, I.; Zheng, J.; James, H. P.; Jin, R.; Baumgart, T. Multivalent Interactions between Molecular Components Involved in Fast Endophilin Mediated Endocytosis Drive Protein Phase Separation. Nat. Commun. 2022, 13 (1), 5017. 10.1038/s41467-022-32529-0

(51) Martin, E. W.; Harmon, T. S.; Hopkins, J. B.; Chakravarthy, S.; Incicco, J. J.; Schuck, P.; Soranno, A.; Mittag, T. A Multi-Step Nucleation Process Determines the Kinetics of Prion-like Domain Phase Separation. Nat. Commun. 2021, 12 (1), 4513. 10.1038/s41467-021-24727-z

(52) Balu, R.; Wanasingha, N.; Mata, J. P.; Rekas, A.; Barrett, S.; Dumsday, G.; Thornton, A. W.; Hill, A. J.; Roy Choudhury, N.; Dutta, N. K. Crowder-Directed Interactions and Conformational Dynamics in Multistimuli-Responsive Intrinsically Disordered Protein. Sci. Adv. 2022, 8 (51), eabq2202. 10.1126/sciadv.abq2202

(53) Hammouda, B. Analysis of the Beaucage Model. J. Appl. Crystallogr. 2010, 43 (6), 1474– 1478. 10.1107/S0021889810033856

(54) Hochmair, J.; Exner, C.; Franck, M.; Dominguez-Baquero, A.; Diez, L.; Brognaro, H.; Kraushar, M. L.; Mielke, T.; Radbruch, H.; Kaniyappan, S.; Falke, S.; Mandelkow, E.; Betzel, C.; Wegmann, S. Molecular Crowding and RNA Synergize to Promote Phase Separation, Microtubule Interaction, and Seeding of Tau Condensates. EMBO J. 2022, 41 (11), EMBJ2021108882. 10.15252/embj.2021108882

(55) Lifshitz, I. M.; Slyozov, V. V. The Kinetics of Precipitation from Supersaturated Solid Solutions. J. Phys. Chem. Solids 1961, 19 (1–2), 35–50. 10.1016/0022-3697(61)90054-3

(56) Tadros, T. Ostwald Ripening. In *Encyclopedia of Colloid and Interface Science*; Tadros, T., Ed.; Springer Berlin Heidelberg: Berlin, Heidelberg, 2013; pp 820–820. 10.1007/978-3-642-20665-8_124

(57) Wagner, C. Theorie Der Alterung von Niederschlägen Durch Umlösen (Ostwald-Reifung). Z. Für Elektrochem. Berichte Bunsenges. Für Phys. Chem. 1961, 65 (7–8), 581–591. 10.1002/bbpc.19610650704

(58) Canchi, D. R.; Jayasimha, P.; Rau, D. C.; Makhatadze, G. I.; Garcia, A. E. Molecular Mechanism for the Preferential Exclusion of TMAO from Protein Surfaces. J. Phys. Chem. B 2012, 116 (40), 12095–12104. 10.1021/jp304298c

(59) Courtenay, E. S.; Capp, M. W.; Anderson, C. F.; Record, M. T. Vapor Pressure Osmometry Studies of Osmolyte−Protein Interactions: Implications for the Action of Osmoprotectants in Vivo and for the Interpretation of “Osmotic Stress” Experiments in Vitro. Biochemistry 2000, 39 (15), 4455–4471. 10.1021/bi992887l

(60) Hernández-Vega, A.; Braun, M.; Scharrel, L.; Jahnel, M.; Wegmann, S.; Hyman, B. T.; Alberti, S.; Diez, S.; Hyman, A. A. Local Nucleation of Microtubule Bundles through Tubulin Concentration into a Condensed Tau Phase. Cell Rep. 2017, 20 (10), 2304–2312. 10.1016/j.celrep.2017.08.042

(61) Das, S.; Lin, Y.-H.; Vernon, R. M.; Forman-Kay, J. D.; Chan, H. S. Comparative Roles of Charge, *π*, and Hydrophobic Interactions in Sequence-Dependent Phase Separation of Intrinsically Disordered Proteins. Proc. Natl. Acad. Sci. 2020, 117 (46), 28795–28805. 10.1073/pnas.2008122117

(62) Dignon, G. L.; Best, R. B.; Mittal, J. Biomolecular Phase Separation: From Molecular Driving Forces to Macroscopic Properties. Annu. Rev. Phys. Chem. 2020, 71 (1), 53–75. 10.1146/annurev-physchem-071819-113553

(63) Lin, Y.; Currie, S. L.; Rosen, M. K. Intrinsically Disordered Sequences Enable Modulation of Protein Phase Separation through Distributed Tyrosine Motifs. J. Biol. Chem. 2017, 292 (46), 19110–19120. 10.1074/jbc.M117.800466

(64) Dignon, G. L.; Zheng, W.; Kim, Y. C.; Best, R. B.; Mittal, J. Sequence Determinants of Protein Phase Behavior from a Coarse-Grained Model. PLOS Comput. Biol. 2018, 14 (1), e1005941. 10.1371/journal.pcbi.1005941

(65) Martin, E. W.; Mittag, T. Relationship of Sequence and Phase Separation in Protein Low-Complexity Regions. Biochemistry 2018, 57 (17), 2478–2487. 10.1021/acs.biochem.8b00008

(66) Phan, T. M.; Kim, Y. C.; Debelouchina, G. T.; Mittal, J. Interplay between Charge Distribution and DNA in Shaping HP1 Paralog Phase Separation and Localization. eLife 2024, 12, RP90820. 10.7554/eLife.90820.3

(67) Schiano Lomoriello, I.; Sigismund, S.; Day, K. J. Biophysics of Endocytic Vesicle Formation: A Focus on Liquid–Liquid Phase Separation. Curr. Opin. Cell Biol. 2022, 75, 102068. 10.1016/j.ceb.2022.02.002

(68) Speicher, T. L.; Li, P. Z.; Wallace, I. S. Phosphoregulation of the Plant Cellulose Synthase Complex and Cellulose Synthase-Like Proteins. Plants 2018, 7 (3), 52. 10.3390/plants7030052

