## additional methods and results for "Characterization of Liquid-Liquid Phase Separation of Companion of Cellulose Synthases under Stress Mimicking Conditions"

### Phase separation enhances the function of salt stress related plant microtubule-associated proteins

*This manuscript has been authored by UT-Battelle, LLC under Contract No. DE-AC05-00OR22725 with the U.S. Department of Energy. The United States Government retains and the publisher, by accepting the article for publication, acknowledges that the United States Government retains a non-exclusive, paid-up, irrevocable, world-wide license to publish or reproduce the published form of this manuscript, or allow others to do so, for United States Government purposes. The Department of Energy will provide public access to these results of federally sponsored research in accordance with the DOE Public Access Plan (http://energy.gov/downloads/doe-public-access-plan)*

**S1. Expression and purification of the N-terminal domains of CC1 and CC2**

The synthetic genes encoding CC1NTD (1-120 aa) and CC2NTD (1-98 aa) were cloned into pET28a+ vector which has a hexahistidine tag at the C-terminal end (GenScript, USA). Both plasmids were transformed into *E. coli* BL21(DE3) competent cells based on previously described constructs (Kesten et al., 2019). Protein overexpression was carried out in Lysogeny broth (LB) with Kanamycin (50 µg/mL) at 37 ºC. Isopropyl-b-D-thiogalactopyranoside (IPTG) was added to a final concentration of 1 mM to induce overexpression when the optical density of the culture, measured at 600nm ($OD_{600}$), reached 0.6 – 0.8. Following the induction, the cell growth was continued for 4h at 37ºC. The cell pellet was collected by centrifugation at 5,700 x g for 30min. The cells were either stored at -80ºC for later use or immediately lysed for protein purification.

Cell pellets were resuspended in the lysis buffer (50mM Tris, 150mM NaCl, cOmplete EDTA-free protease inhibitor cocktail (Roche), pH 8) and lysed using a Branson 450 Digital Sonifier at 60% amplitude for 5min (pulse on for 2s and off for 10 s) in an ice bath. The lysed cells were centrifuged at 17,000 x g for 20min and the clarified lysate was filtered using 0.45µm filter. The recombinant CC N-terminal domains were purified using a HisTrap™ FF (Cytiva Life Sciences, USA) column and an ÄKTA Start Chromatography system (GE Healthcare Life Sciences, USA). The purification of CC1NTD is described elsewhere (Gurumoorthy et al., 2025). Nickel-nitriloacetic affinity chromatography (NiNTA) was used to purify the histidine-tagged CC2NTD. The column was equilibrated with 5 CVs buffer A (50mM Tris-HCl, 200mM NaCl, 20mM imidazole, pH 8) followed by loading the clarified lysate at a flow rate of 1 ml/min. The bound protein was first washed with 10 CVs of a mixture of 96% buffer A and 4% buffer B (50mM Tris-HCl, 200mM NaCl, 250mM imidazole, pH 8) and was then eluted with a linear gradient of 4 – 100% of buffer B (10 CVs). CC2NTD eluted from the column in 70% buffer B. The resin was further washed with 3 CVs of 100% buffer B to remove any residual bound protein. CC2NTD was concentrated by centrifugation using Amicon centrifugal devices with the molecular weight cutoff of 3000 Da. The concentrated protein was injected into a 2mL loop connected to a Superdex 200 Increase 10/300 GL column (GE Healthcare, USA) that was equilibrated with 2 CVs of gel filtration buffer (50mM Tris-HCl, 150mM NaCl, 2mM TCEP, pH 8.0). The protein was eluted with 1 CV gel filtration buffer, and 0.5mL fractions were collected for further analysis. The fractions from SEC that contained pure CC2NTD were either flash frozen using liquid nitrogen and stored at -80°C until later use or immediately used for characterization studies.

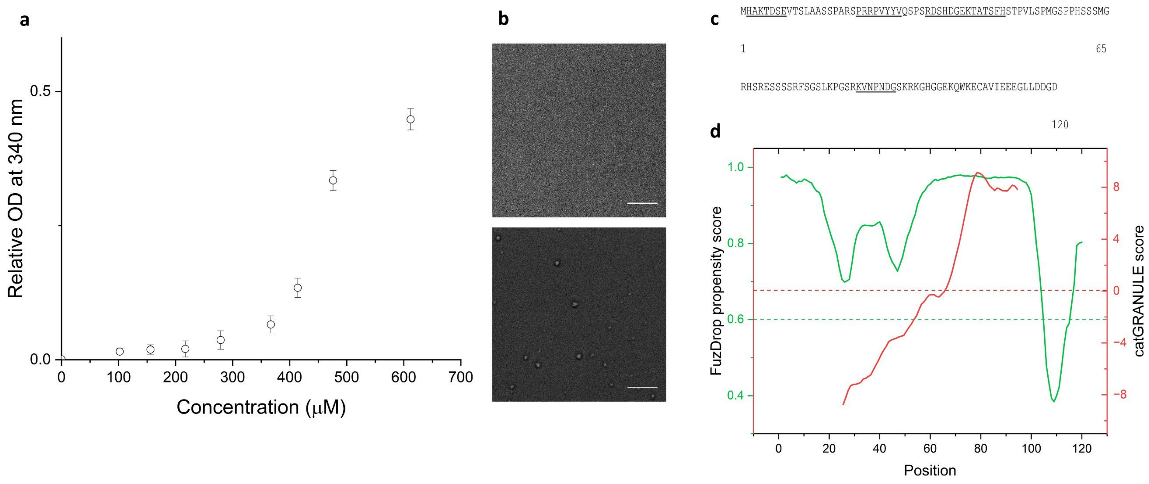


*Figure S1. CC1NTD undergoes LLPS under non-reducing conditions (a) CC1NTD turbidity measured using UV-visible spectrophotometry at 340nm at different concentrations; (b) DIC images of (top) CC1NTD (0μM) and (bottom) CC1NTD (450μM). Scale bar – 10μm (c) CC1NTD amino acid sequence was analyzed with FuzDrop. The prediction of aggregation-prone regions is underlined; (d) Plot of CC1NTD phase separation propensity vs. amino acid position predicted by FuzDrop algorithm (green line) and catGRANULE (red line). Dashed green line and dashed red line indicate the cutoff above which lies the positive prediction for LLPS propensity.*

#### S2. Fluorescent dye labeling of CC1 and CC2 N-terminal domains

The amino-terminal $NH_2$ group of both CC1NTD and CC2NTD was covalently labeled using Alexa Fluor 488 NHS succinimidyl ester (Invitrogen, USA) for fluorescent imaging and assays. First, proteins (~ 2 mg/ml) were buffer exchanged to 100 mM sodium bicarbonate buffer (pH 8.3) by dialysis using Slide-A-Lyzer™ 0.5mL cassette (3.5 kDa molecular weight cutoff) for 16h at 4°C. Then, an approximately 10-fold molar excess of dye dissolved in dimethylformamide was added drop-by-drop to 100 μL of protein (2 mg/mL) while mixing with a Thermomixer® (Eppendorf, USA). The conjugation of protein and dye was continued at room temperature with shaking at 350rpm for 1 h. Size exclusion chromatography was used to separate the unreacted dye from the protein. A Superdex Increase 75 5/150 GL column (Cytiva Life Sciences, USA) was equilibrated with 2 CVs of 20mM HEPES, 150mM NaCl (pH 8.0) buffer before applying the reaction mixture using a 2mL loop. The labeled CC1NTD and CC2NTD were eluted using an isocratic gradient of 1.5 CVs buffer at a flow rate of 0.5 mL/min. The protein concentration and the degree of labeling were determined using the manufacturer's protocol, as described below. The concentration of the conjugated protein (Alexa488 conjugated protein) was determined as follows:

$$[Alexa488\ conjugated\ protein] = \frac{A_{280} - (A_{495} * CF)}{\varepsilon} \qquad (S1)$$

where $A_{280}$ and $A_{495}$ are the absorbance values at 280 and 495nm; CF is the correction factor of dye (0.11 for Alexa488 NHS); and ε is the molar extinction coefficient of the protein. The degree of labeling (DOL) was determined as follows:

$$DOL = \frac{A_{495} * MW}{[Alexa488\ conjugated\ protein] * \varepsilon_{dye}} \qquad (S2)$$

where MW is the molecular weight of the protein and $\varepsilon_{dye}$ is the dye molar extinction coefficient of 71,000 $cm^{-1}M^{-1}$ at the maximum absorbance wavelength $\lambda_{max}$ = 495 nm. The labeled CC1NTD and CC2NTD were immediately flash frozen and stored in -80°C until further use.

**S3. Sample preparation for microscopy**

For fluorescence imaging and FRAP experiments, unlabeled CC1NTD or CC2NTD was mixed with Alexa Fluor 488–labeled CC1NTD or CC2NTD, respectively, at a labeling ratio of ~4% (mol/mol). Protein mixtures were combined with either Tris buffer (control) or TMAO-containing buffer, placed onto 35 mm glass-bottom dishes (Nunc™, Thermo Fisher Scientific, USA), and imaged immediately. For tubulin co-condensation and FRAP experiments, HiLyte 647-labeled tubulin was mixed with unlabeled tubulin at a 1:10 molar ratio in general tubulin buffer (Cytoskeleton Inc., USA). The final concentration of tubulin in the condensates was maintained at 54μM. Preformed CC1NTD condensates were then added to the tubulin mixture, transferred to glass-bottom dishes, and imaged immediately. To induce microtubule polymerization during imaging, ice-cold microtubule polymerization buffer was added directly to the condensate–tubulin mixture prior to imaging. For CC1NTD-CC2NTD co-condensation experiments, CC1NTD condensates were first formed and then mixed with Alexa Fluor 488-labeled CC2NTD at a 4:1 (v/v) ratio and imaged immediately. For experiments involving tubulin, the labeled/unlabeled tubulin mixture described above was added to preformed CC1NTD-CC2NTD condensates prior to imaging. The apparent partition coefficient was calculated as the ratio of the fluorescence intensity within the condensates to that in the surrounding phase. Droplet intensity and size were quantified using ImageJ (Schneider et al., 2012). For each condition, at least 10 droplets from three independent replicates were analyzed. Data are reported as mean ± SEM. Imaging was performed using a Leica SP8 confocal microscope (Leica Microsystems, Germany) equipped with a 63× oil immersion objective.

#### S4. Dynamic Light Scattering Analysis

All measurements were collected for 30 min with 10 s acquisition intervals for a total of 60 acquisitions. DYNAMICS® v 7.0 was used to analyze experimental data. All autocorrelation functions were averaged and fitted using an in-built nonlinear least-squares fitting algorithm. The hydrodynamic radii of the samples were estimated by fitting autocorrelation functions of scattered light intensity to a Rayleigh sphere model assuming sphere-like condensate formation under osmolyte conditions. The hydrodynamic radius, $R_0$, was calculated using the Stokes-Einstein equation as follows:

$$R_0 = \frac{\kappa_B T}{6\pi\eta D} \quad (S3)$$

where $\kappa_B$is the Boltzmann constant, T is the temperature, $\eta$ is the viscosity, and $D$ is the rate of diffusion. For time-resolved DLS (trDLS) measurements, the growth of spherical particles was fitted using the following power law [34]:

$$R_0 = a * t^b \quad (S4)$$

where $a$ and $b$ are fitting parameters, and t is the time in s. The coarsening exponent, $b$, indicates the rate at which the phase transition occurs.

#### S5. Fluorescence recovery after photobleaching analysis

FRAP data analysis was performed as described previously [42]. The data were fit to the following exponential function:

$$y = y0 + A\left(1 - e^{-\frac{x}{\tau}}\right) \quad (S5)$$

where $y0$ is the offset; $A$ is the amplitude; and the half-time recovery rate ($t_{\frac{1}{2}}$) is calculated using $t_{\frac{1}{2}} = 0.69 * \tau$. The apparent diffusion coefficient $D_{app}$ was estimated after measuring the radius of the bleached area ($r$) using the equation:

$$D_{app} = \frac{r^2}{\tau} \qquad (S6)$$

#### S6. Small-angle X-ray scattering analysis of CC1NTD in solution

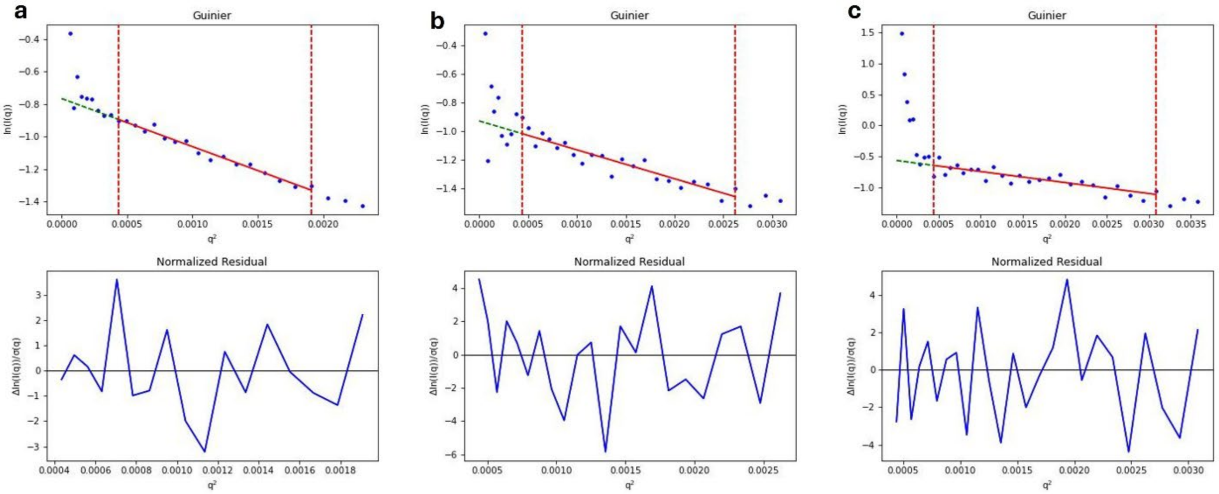


*Figure S2. Guinier analysis of CC1NTD at different TMAO concentrations. Guinier and normalized residuals of CC1NTD at (a) 0 M TMAO; (b) 0.75M TMAO; and (c) 1.1M TMAO.*

#### S7. Small-angle neutron scattering data analysis

The SANS profiles were analyzed using a combined Unified fit (Beaucage, 1996, 1995) and core–shell cylinder model. The Unified Fit contribution describes the characteristic size and scaling behavior of mesoscale organization within the condensates, whereas the core–shell cylinder term describes the cross-section of smaller elongated assemblies that are visible in solutions containing tubulin. The total scattering intensity was therefore expressed as the sum of the two contributions,

The total scattering intensity was expressed as:

$$I(Q) = I_{\mathrm{UF}}(Q) + I_{\mathrm{CS}}(Q) + I_{\mathrm{bkg}} \qquad (S7)$$

where $I_{\mathrm{UF}}(Q)$ is the Unified Fit contribution, $I_{\mathrm{CS}}(Q)$ is the core–shell cylinder contribution, and $I_{\mathrm{bkg}}$ is a $Q$-independent background. Additional details on the fitting approach and equations are provided in the Supplementary Information.

For a single structural level, the Unified contribution was defined as

$$I_{\mathrm{UF}}(Q) = G \exp\left(-\frac{Q^2 R_g^2}{3}\right) + B\left[\frac{\mathrm{erf}^3\left(QR_g/\sqrt{6}\right)}{Q}\right]^P \qquad (S8)$$

where $G$ is the Guinier prefactor, $R_g$ is the radius of gyration, $B$ is the power-law prefactor, and $P$ is the power-law exponent. The first term describes the Guinier regime associated with the characteristic size of the structures, whereas the second term describes their power-law scattering. In this study, the $R_g$values were calculated for the low-$Q$ regime in the data, representing the mesoscale organization of the condensates. The values obtained likely represent lower boundary for the size of the particles observed rather than the size of the proteins or tubulin assemblies as the SANS data did not fully capture a Guinier regime. The power-law exponent, $P$, describes how scattering intensity changes with $Q$ in the scaling regime. For $1 < P < 3$, $P$ can be interpreted as an apparent mass-fractal dimension, ($D_m$), with lower values indicating a more open and weakly space-filling organization. For ($3 < P < 4$), the scattering is consistent with surface-fractal or interface-dominated behavior, and an apparent surface-fractal dimension can be estimated as ($Ds = 6 - P$).

The core–shell cylinder model describes a homogeneous cylindrical core with radius $R$, length $L$, and scattering length density $\rho_c$, surrounded by a shell of uniform thickness $T$ and scattering length density $\rho_s$. The overall particle radius is $R + T$ and an overall diameter is $2(R + T)$. The particle is dispersed in a solvent with scattering length density $\rho_{solv}$.

The core–shell cylinder intensity was calculated from the orientationally averaged squared scattering amplitude:

$$I_{\mathrm{CS}}(Q) = \frac{\mathrm{Scale}}{V_t} \int_0^{\pi/2} |F(Q,\alpha)|^2 \sin\alpha \; d\alpha \qquad (S9)$$

where α is the angle between the cylinder axis and the scattering vector $Q$, $V_t$ is the total particle volume, and $Scale$ is proportional to the particle volume fraction for data measured on an absolute intensity scale.

The total scattering amplitude is

$$F(Q,\alpha) = F_{core}(Q,\alpha) + F_{outer}(Q,\alpha) \qquad (S10)$$

where the core contribution is

$$F_{\mathrm{core}}(Q,\alpha) = (\rho_c - \rho_s)V_c A_{\mathrm{cyl}}(Q,\alpha; R, L) \qquad (S11)$$

and the contribution from the complete core–shell cylinder is

$$F_{outer}(Q,\alpha) = (\rho_s - \rho_{solv})V_t A_{cyl}(Q,\alpha; R + T, L + 2T) \qquad (S12)$$

The normalized scattering amplitude of a homogeneous cylinder is

$$A_{\mathrm{cyl}}(Q, \alpha; R, L) = \frac{\sin(QL\cos\alpha/2)}{QL\cos\alpha/2} \frac{2J_1(QR\sin\alpha)}{QR\sin\alpha} \tag{S13}$$

where $J_1$ is the first-order Bessel function. The sine term describes scattering along the cylinder axis, whereas the Bessel-function term describes scattering across its circular cross-section.

The cylindrical core volume is:

$$V_c = \pi R^2 L \tag{S14}$$

and the total volume of the core–shell cylinder is:

$$V_t = \pi(R + T)^2(L + 2T) \tag{S15}$$

The SLD differences $\rho_c - \rho_s$ and $\rho_s - \rho_{solv}$ represent the scattering contrasts at the core–shell and shell–solvent interfaces, respectively (Table S4).

Combining both structural contributions, the complete fitting function was

$$I(Q) = G\exp\left(-\frac{Q^2R_g^2}{3}\right) + B\left[\frac{\mathrm{erf}^3\left(QR_g/\sqrt{6}\right)}{Q}\right]^P + \frac{\mathrm{Scale}}{V_t}\int_0^{\frac{\pi}{2}} |F(Q,\alpha)|^2 \sin\alpha \, d\alpha + I_{\mathrm{bkg}} \tag{S16}$$

The cylinder core radius and shell thickness were fitted as global parameters shared across datasets. Core and shell SLDs were fitted separately for each contrast-defined sample and were shared between the corresponding long- and short-configuration measurements. The cylinder length was fixed at 800 nm and was not fitted. This value was chosen to represent a structure substantially longer than its cross-sectional dimensions, so the fit primarily reports the cross-section rather than the true contour length. The cylinder scale and the Unified parameters $R_g$, power-law exponent $P$, $B$, and $G$ , and background were fitted independently for each dataset. The solvent SLD and cylinder length were fixed at their specified initial values. The reported uncertainties correspond to the one-standard-deviation estimates obtained from the DREAM posterior distributions.

**Table S1. Core–shell cylinder parameters**

| **Core-Shell cylinder structural parameters** | **Value (nm)** |
|---|---|
| Core radius, R | 8.4±0.1 nm |
| Core diameter, 2R | 16.7±0.3 nm |
| Shell thickness, T | 5.8±0.3 nm |
| Outer diameter, 2(R+T) | 28.4±0.6 nm |
| Core cylinder length, L (Fixed) | 800 nm |
| SLD core, 85% $D_2O$ hCC1NTD: Tubulin | 5.5±0.1 Å$^{-2}$ |
| SLD shell, 85% $D_2O$ hCC1NTD: Tubulin | 2.8±0.3 Å$^{-2}$ |
| SLD core, 85% $D_2O$ dCC1NTD: Tubulin | 5.4 ± 0.1 Å$^{-2}$ |
| SLD shell, 85% $D_2O$ dCC1NTD: Tubulin | 4.4 ± 0.1 Å$^{-2}$ |
| SLD core, 42% $D_2O$ dCC1NTD: Tubulin | 1.8 ± 0.3 Å$^{-2}$ |
| SLD shell, 42% $D_2O$ dCC1NTD: Tubulin | 3.2 ± 0.1 Å$^{-2}$ |
| SLD buffer 85% $D_2O$ | 4.98 Å$^{-2}$ |
| SLD buffer 42% $D_2O$ | 2.97 Å$^{-2}$ |

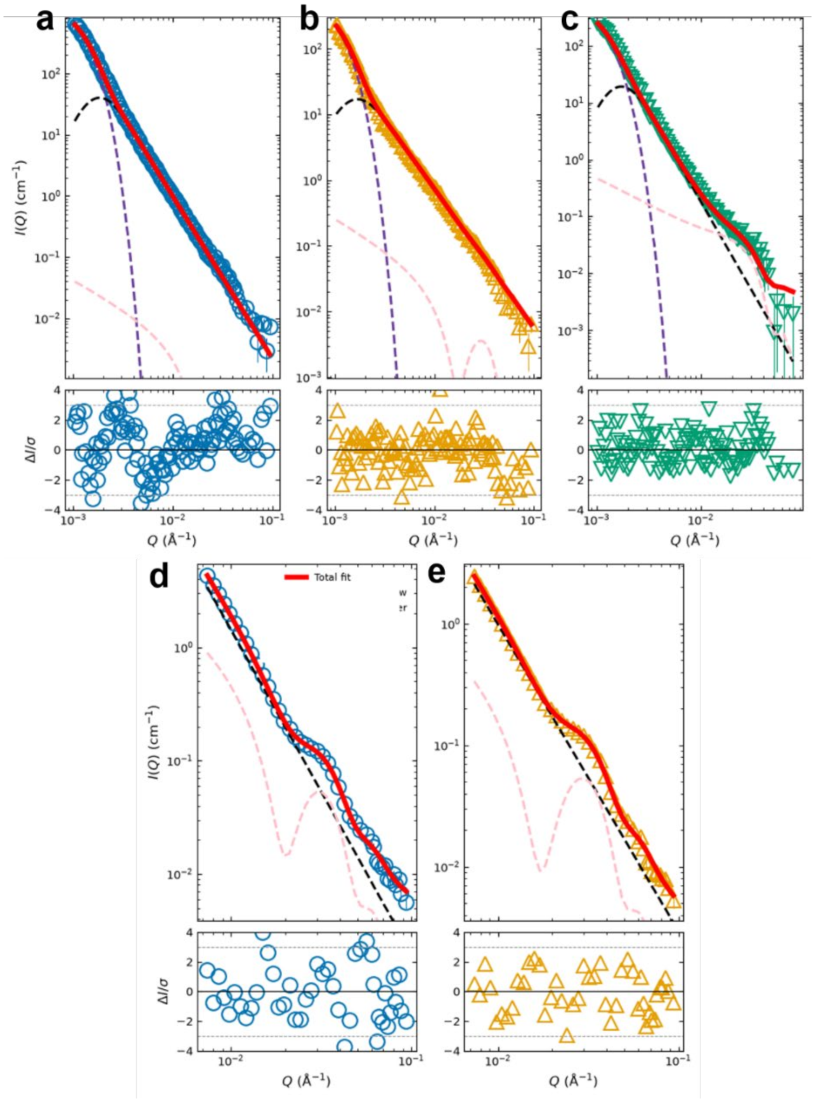


*Figure S3. Contrast-variation SANS analysis of CC1NTD–tubulin condensates***.** *Panels a – c show SANS profiles and corresponding model fits are shown for measurements collected using the long instrument configuration. (a) fully protiated CC1NTD–tubulin condensates in 85% $D_2O$ (open blue circles); (b) deuterated CC1NTD–tubulin condensates in 85% $D_2O$ (yellow triangles); and (c) deuterated CC1NTD–tubulin condensates in 42% $D_2O$ (green inverted triangles). The corresponding short-configuration measurements are shown for fully protiated CC1NTD–tubulin condensates in 85% $D_2O$ (d) and dCC1NTD–tubulin condensates in 85% $D_2O$ (e), using the same symbols and colors as in panels a and b, respectively. Error bars indicate uncertainties arising from neutron counting statistics. The combined Unified Fit and core–shell cylinder model is shown as a solid red line. The Guinier and power-law contributions of the Unified Fit are shown as dashed purple and black lines, respectively, while the core–shell cylinder contribution is shown as a dashed pink line. Normalized residuals are plotted beneath each scattering profile and colored the same as the corresponding experimental data. The cylinder core radius and shell thickness were treated as global parameters shared across all datasets, whereas core and shell SLDs were fitted separately for each contrast condition and shared between corresponding instrument configurations and are present in the* ***Table S1****.*

#### S8. Bioinformatics and physicochemical analysis of CC2 N-terminal domain

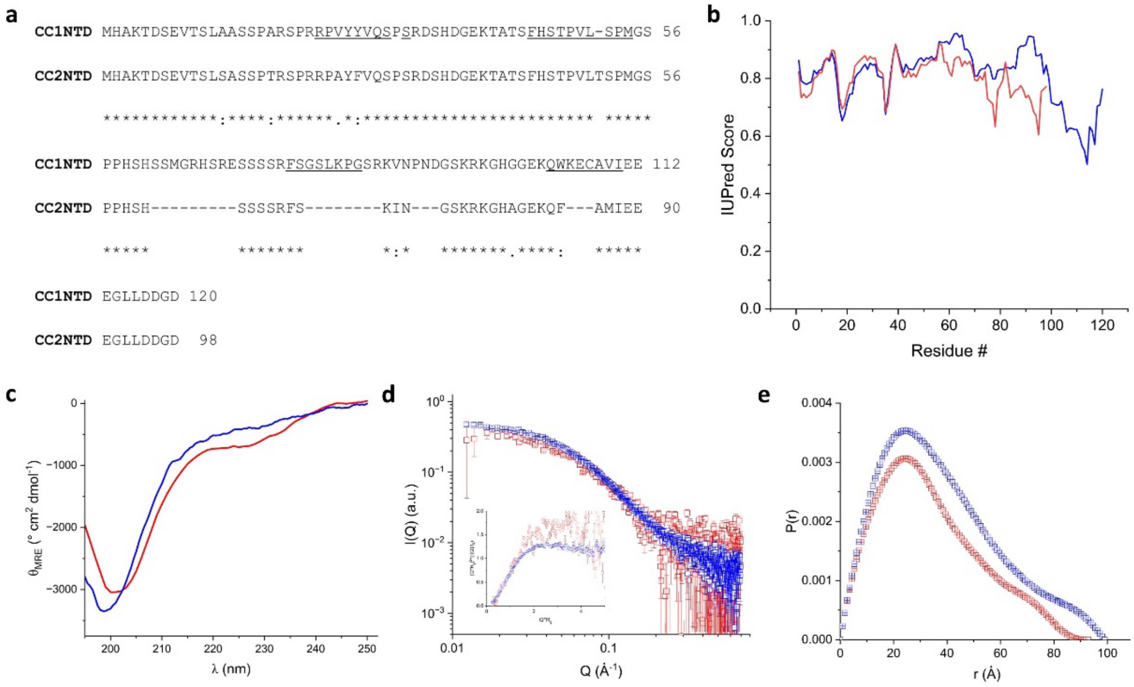


*Figure S4. Comparison of properties of CC1NTD and CC2NTD (a) sequence alignment of CC1NTD (120 aa) and CC2NTD (98 aa) showing sequence homology between the proteins. NMR identified microtubule binding regions of CC1NTD are underlined; (b) Disorder prediction of CC1NTD (blue line) and CC2NTD (red line) using IUPred2 shows that both proteins are intrinsically disordered; (c) CD spectrum of CC2NTD (red line) compared with CC1NTD (blue line) shown as molar ellipticity vs. wavelength shows high negative ellipticity at 200nm indicating CC2NTD is intrinsically disordered like CC1NTD; (d) SAXS of 60μM CC2NTD and CC1NTD shown as measured intensity vs. scattering vector Q. Inset – Kratky representation of CC2NTD and CC1NTD; (e) Pairwise distribution functions of CC2NTD and CC1NTD shows that it has an elongated conformation as evidenced from an extended tail. CC1NTD and CC2NTD are shown as open blue squares and open red squares, respectively.*

*Table S2. Comparison of secondary structure content of CC2NTD and CC1NTD using BestSel (Micsonai et al., 2018)*

| Secondary structure | CC2NTD | CC1NTD |
|---|---|---|
| Helix | 1 | 1 |
| Sheet | 32 | 18 |
| Others | 67 | 81 |

*Table S3. Structural parameters from SAXS of CC2NTD compared to CC1NTD*

| **Parameter** | **CC2NTD** | **CC1NTD** |
|---|---|---|
| $R_g$ (Guinier) | 29.4 ± 1.2 Å | 28.9 ± 0.5 Å |
| $I_0$ (Guinier) | 0.37 ± 0.01 | 0.47 ± 0.01 |
| $R_g$ (GNOM) | 27.7 ± 0.45 Å | 30.8 ± 0.18 Å |
| $D_{max}$ | 95 Å | 99 Å |
| $I_0$ (GNOM) | 0.35 ± 0.01 | 0.45 ± 0.01 |
| Theoretical molecular weight | 12.9 kDa | 14.8 kDa |
| Experimental molecular weight (Bayes) | 16.8 kDa | 13.6 kDa |

**S9. Microtubule binding assay**

A microtubule-binding spin-down assay using ultracentrifugation was performed as described in the manufacturer's protocol (Cytoskeleton, Inc, USA). Porcine brain tubulin (T240 Cytoskeleton, Inc, USA) was dissolved in the general tubulin buffer (80mM PIPES [piperazine-N,N′-bis(2-ethanesulfonic acid)], 2mM $MgCl_2$, 0.5mM EGTA pH 6.9) plus 1mM GTP and 5%glycerol, flash frozen in liquid nitrogen and stored as 20μL aliquots (5 mg/mL) at -80°C until used. For microtubule preparation, the aliquots were quickly thawed and polymerized in the presence of cushion buffer (80mM piperazine-N,N′-bis(2-ethanesulfonic acid), 2mM $MgCl_2$, 0.5mM EGTA, 60% glycerol pH 6.9) and stabilized using 2mM paclitaxel. This yielded 0.5μM of tubulin dimer-containing microtubules. Variable concentrations of proteins (0 – 30μM) were added to a fixed microtubule concentration (0.5μM tubulin) and made up to a final volume of 50 μL using general tubulin buffer. The reaction mixture was incubated for 30min at room temperature. To each reaction tube, 100μL of cushion buffer with 2 mM paclitaxel were then added. The mixture was spun down by ultracentrifugation (Sorvall™ Thermo Fisher Scientific, USA) at 100,000 x g for 40min at 20°C. From each reaction tube, 50 μL supernatant was removed and mixed with Laemmli sample buffer according to the manufacturer's protocol (Cytoskeleton Inc, USA). Similarly, each pellet was resuspended in Laemmli sample buffer for gel electrophoresis. For sodium dodecyl sulfate polyacrylamide gel electrophoresis (SDS PAGE), samples containing Laemmli buffer were heated to 95°C for 5min, briefly spun, and loaded on to 4-20% Mini-PROTEAN® TGX Stain-Free™ gels (Bio Rad, USA). The electrophoresis was conducted using Mini-PROTEAN Tetra gel electrophoresis unit (Bio Rad, USA) at constant 300V for 20min. The gels were then stained with GelCode™ blue stain (Thermo Fisher Scientific, USA) for 1h by gentle rocking, followed by overnight destaining in deionized water. The gel images were analyzed for band intensity quantification using ImageJ software. The concentration of protein and relative intensity quantified from SDS-PAGE of pellets (protein bound to tubulin) were plotted. Experiments were conducted in three replicates. Data were presented as mean ± sem. The dissociation constant was calculated by fitting the data using OriginLab to Michaelis-Menten equation as follows:

$$y = \frac{V_{max} * x}{K_M + x} \tag{S17}$$

where $V_{max}$ is the maximum velocity at and $K_M$is the dissociation constant.

#### S10. Negative stain electron microscopy of CC2NTD-microtubule interactions

Transmission electron microscopy was performed using a JEOL JEM 1400-Flash transmission electron microscope (JEOL, Japan) operated at 80 kV. First, the carbon-coated side of the hexagonal mesh copper grids (Ted Pella, Redding, CA, USA) was placed on the samples of microtubules or a CC2NTD-microtubules mixture (10μL) for a minute. Then the excess of sample was removed by placing the grid in water for 5 s. Subsequently, the grids were stained using uranyl-less solution for 1.5min. After staining, excess stain was removed by blotting on to a Whatman filter and the grids were dried overnight at room temperature before viewing under microscope.

#### S11. SAXS Analysis of CC2-microtubule interactions

The structure factor for CC-induced microtubule bundles was fitted to Lorentzian peaks using the following equation:

$$S(q) = c + \frac{2A}{\pi} \times \frac{w}{4(Q - Q_{hk})^2 + w^2} \qquad (S18)$$

where $Q_{hk} = Q_{10} \times \sqrt{h^2 + k^2 + hk}$ , A is the area, w is full width at half maximum, and c is offset. Here hk indices refer to 2D hexagonal lattices corresponding to 10, 11, 20, 21, 30, 22, and so on, and $Q_{10}$ is the first peak from hexagonal lattice. The lattice parameter ($a_H$) of bundled microtubules [41] is given by the following equation:

$$a_H = \frac{4\pi}{\sqrt{3} \times Q_{10}} \qquad (S19)$$

**S12. Characterization of CC2NTD-microtubule complex**

For functional characterization of CC2NTD, centrifugation-based microtubule pull-down assay was performed to obtain binding kinetics of CC2NTD and microtubule interactions. The assay yielded a dissociation constant of 11.2 ± 2μM comparable to previously reported values for CC1NTD; 10.5 ± 2μM (Gurumoorthy et al., 2025) and 9.6 ± 2μM (Endler et al., 2015). The SAXS profile of the microtubule suspension after the addition of CC2NTD showed the presence of peaks indicating a regular arrangement of the scattering particles, which is clearly different from the microtubules alone. A Lorentzian function structure factor (Eq.S18) was used to fit the data and yielded peak distances that are consistent with cylinders arranged as 2D hexagonal array with a spatial relationship, $Q_{hk}/Q_{10}$, of 1: √3 : √4 : √7 : √9 (Needleman et al., 2004). However, the primary hexagonal peak $Q_{10}$ was not as prominent as in the CC1NTD-induced microtubule bundles and the estimated microtubule center-to-center distance ($a_H$, Eq. S7) was ~ 380Å. This value was somewhat larger than $a_H$ for CC1NTD, which was 325 Å (Gurumoorthy et al., 2025), indicating tighter microtubule bundling by CC1NTD as compared with CC2NTD. CC2NTD is shorter than CC1NTD and lacks residues that were identified to interact with microtubules (Kesten et al., 2019), as described above, which may explain why CC2NTD may not be as efficient in bundling microtubules. However, it is clear that CC1NTD and CC2NTD play functionally similar roles.

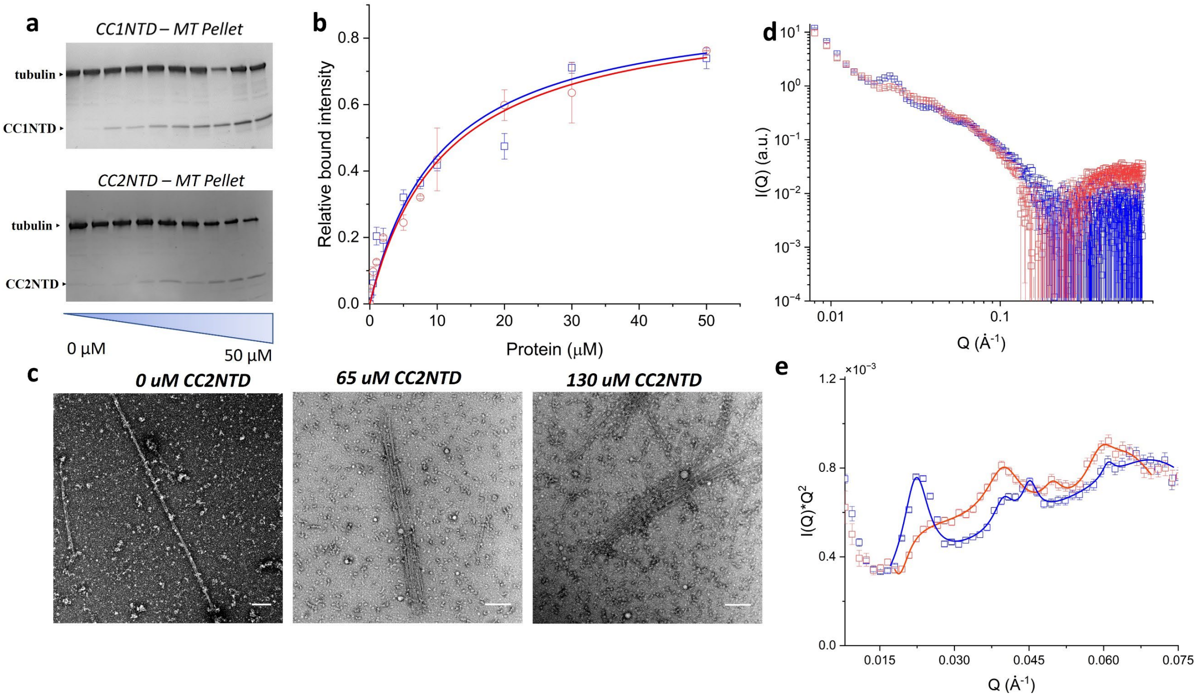


*Figure S5. Comparison of CC1NTD and CC2NTD interactions with microtubules (a) Microtubule binding assays of CC1NTD (top) and CC2NTD (bottom) shown as SDS-PAGE analysis of pellets collected from centrifuged samples of fixed concentration of microtubules (2µM tubulin dimer) incubated with increasing concentrations of protein (0 – 50µM); (b) Microtubule binding assay results plotted as relative bound intensity of protein determined from SDS-PAGE vs. concentration of incubated protein (n = 3); (c) TEM images of free microtubules and CC2NTD-bundled microtubules. showing the concentration dependence of CC2NTD on the microtubule bundle thickness. Scale bar – 2µm; (d) SAXS of CC1NTD (100µM) (open blue squares) and CC2NTD (120µM) (open red squares) bundle microtubules (tubulin dimer 18µM); (e) Kratky representation of CC1NTD and CC2NTD SAXS data highlighting the peak positions for microtubule bundles and hexagonal Lorentzian structure factor peak fits shown as blue and red lines, respectively.*

*Table S4. Fitting parameters from SAXS of CC1NTD and CC2NTD microtubule bundles*

| CC1NTD microtubules | | | CC2NTD microtubules | | |
|---|---|---|---|---|---|
| **Peak Center $Q_{hk}$ ($Å^{-1}$)** | **Full Width at Half Maximum ($Å^{-1}$)** | **Peak Area ($Å^2$)** | **Peak Center $Q_{hk}$ ($Å^{-1}$)** | **Full Width at Half Maximum ($Å^{-1}$)** | **Peak Area ($Å^2$)** |
| 0.022 | 0.005 ± 0.01 | $4.07 \times 10^{-13}$ | 0.019 | 0.005 ± 0.01 | $1.8 \times 10^{-7}$ |
| 0.04 | 0.007 ± 0.01 | $1.345 \times 10^{-13}$ | 0.04 | 0.008 ± 0.01 | $3.3 \times 10^{-6}$ |
| 0.045 | 0.023 ± 0.01 | $1.97 \times 10^{-10}$ | 0.049 | 0.005 ± 0.01 | $7.3 \times 10^{-7}$ |
| 0.06 | 0.029 ± 0.08 | $9.4 \times 10^{-10}$ | 0.06 | 0.007 ± 0.01 | $1.9 \times 10^{-7}$ |
| 0.065 | 0.025 ± 0.04 | $7.18 \times 10^{-10}$ | 0.065 | 0.015 ± 0.01 | $6.8 \times 10^{-6}$ |

**Movie S1.** Time lapse movie of Alexa Fluor 488 labeled CC1NTD condensates

**Movie S2.** Time lapse movie of CC1NTD condensates in the presence of Alexa Fluor 88 labeled CC2NTD

#### References


Beaucage, G., 1996. Small-Angle Scattering from Polymeric Mass Fractals of Arbitrary Mass-Fractal Dimension. J. Appl. Crystallogr. 29, 134–146. https://doi.org/10.1107/S0021889895011605

Beaucage, G., 1995. Approximations Leading to a Unified Exponential/Power-Law Approach to Small-Angle Scattering. J. Appl. Crystallogr. 28, 717–728. https://doi.org/10.1107/S0021889895005292

Endler, A., Kesten, C., Schneider, R., Zhang, Y., Ivakov, A., Froehlich, A., Funke, N., Persson, S., 2015. A Mechanism for Sustained Cellulose Synthesis during Salt Stress. Cell 162, 1353–1364. https://doi.org/10.1016/j.cell.2015.08.028

Gurumoorthy, V., Hicks, A., Tiruvadi-Krishnan, S., Leite, W.C., Chodankar, S., Kolape, J.L., Smith, J.C., Lamichhane, R., O'Neill, H., 2025. The N-terminal domain of COMPANION OF CELLULOSE SYNTHASE1 promotes microtubule array formation in Arabidopsis. Plant Physiol. 199, kiaf392. https://doi.org/10.1093/plphys/kiaf392

Kesten, C., Wallmann, A., Schneider, R., McFarlane, H.E., Diehl, A., Khan, G.A., Van Rossum, B.-J., Lampugnani, E.R., Szymanski, W.G., Cremer, N., Schmieder, P., Ford, K.L., Seiter, F., Heazlewood, J.L., Sanchez-Rodriguez, C., Oschkinat, H., Persson, S., 2019. The companion of cellulose synthase 1 confers salt tolerance through a Tau-like mechanism in plants. Nat. Commun. 10, 857. https://doi.org/10.1038/s41467-019-08780-3

Micsonai, A., Wien, F., Bulyáki, É., Kun, J., Moussong, É., Lee, Y.-H., Goto, Y., Réfrégiers, M., Kardos, J., 2018. BeStSel: a web server for accurate protein secondary structure prediction and fold recognition from the circular dichroism spectra. Nucleic Acids Res. 46, W315–W322. https://doi.org/10.1093/nar/gky497

Needleman, D.J., Ojeda-Lopez, M.A., Raviv, U., Miller, H.P., Wilson, L., Safinya, C.R., 2004. Higher-order assembly of microtubules by counterions: From hexagonal bundles to living necklaces. Proc. Natl. Acad. Sci. 101, 16099–16103. https://doi.org/10.1073/pnas.0406076101

Schneider, C.A., Rasband, W.S., Eliceiri, K.W., 2012. NIH Image to ImageJ: 25 years of image analysis. Nat. Methods 9, 671–675. https://doi.org/10.1038/nmeth.2089
